# Charge Imbalance Drives Salt-Optimized Nucleosome Phase Separation under Physiological Conditions

**DOI:** 10.64898/2026.07.08.737367

**Authors:** Irene Silvernail, Yang Zhang, Xun Chen, Xingcheng Lin

## Abstract

Liquid-liquid phase separation (LLPS) of chromatin contributes to genome organization and regulates genome accessibility to control gene expression. Despite advances in identifying the environmental conditions that promote chromatin condensation, the specific molecular interactions that initiate condensate formation, as well as the physical mechanisms by which DNA mechanics and epigenetic modifications modulate the resulting interaction network, remain unclear. Here, we utilize a residue-resolution, coarse-grained protein-DNA model to simulate nucleosome interactions across diverse ionic and structural conditions. Our simulations reveal non-monotonic salt-dependent phase-separation behavior, with optimal nucleosome condensation occurring under physiological salt conditions. Such a behavior was caused by the charge-imbalanced polyampholytic nature of nucleosomes, which drives competition between local protein-DNA attractions and global DNA-DNA repulsion. We further demonstrate that the conformational flexibility of nucleosomal DNA promotes unwrapping of DNA from the histone core, thereby strengthening histone-DNA interactions and enhancing condensate formation. Finally, we show that acetylation of histone H3 and H4 tails significantly reduces inter-nucleosomal interactions and increases nucleosome dynamics within condensates. Together, our study establishes a quantitative link between microscopic molecular interactions and macroscopic material properties, providing new insights into how mechanical constraints and epigenetic modifications could cooperatively tune genome architecture.

## Introduction

Accumulating evidence suggested that chromatin phase separation and the formation of the membraneless condensates is a fundamental mechanism contributing to genome organization.^1–11^ Despite extensive studies from Hi-C,^12–16^ imaging,^2,4,17–21^ and theoretical studies,^22–25^ the primary molecular interactions that drive chromatin condensation remain enigmatic.^9^ Recent discoveries support contributions from nucleosomal interactions mediated by multi-valent ions,^6,15,26–29^ regulatory proteins,^3,30–33^ and histone post-translational modifications.^4,5,11^ Among these factors, intrinsic nucleosomal interactions and their modulation by ionic concentrations are particularly attractive as they are directly linked to the fundamental mechanism underpinning genome organization and the material properties of chromatin throughout the cell cycle.^4,5,21,34^

The stability and structural organization of biomolecular condensates are strongly influenced by electrostatic interactions,^35,36^ which, in the context of chromatin, originate from the hybrid polyampholytic nature of nucleosomes. ^37–39^ As noted in recent scaling theories, the conformations of polyampholytic intrinsically disordered proteins (IDPs), such as histone tails, are governed by a delicate balance between short-range attractions and long-range repulsions.^40^ This balance is highly sensitive to the ionic environment: changes in salt concentration can screen electrostatic interactions, promote transitions between collapsed and extended tail states, and thereby modulate the nucleosome’s effective hydrodynamic radius and its ability to engage in intermolecular contacts. Furthermore, even in polyelectrolyte-polymer complex systems, Coulombic forces can drive the assembly of liquid-like coacervates. These processes are governed by factors such as local charge density and polarization-induced attractions, through which positively charged histone tails can effectively bridge the negatively charged DNA of neighboring units.^41^ Understanding these ion-modulated electrostatic contributions at the single-nucleosome level is therefore essential for predicting the macroscopic phase behavior and stability of chromatin condensates.

In addition to intrinsic nucleosomal interactions and their sequence-specific modulation, epigenetic modifications are essential for regulating gene expression in cells that share the same genetic content.^42–44^ One important mechanism by which these modifications regulate gene expression is through modulating the properties of phase-separated chromatin.^4,11^ Multiple experiments have shown that histone acetylation antagonizes chromatin phase separation,^4,11^ aligning with other experiments showing histone acetylations destabilize nucleosomes,^45^ weakens inter-nucleosome interactions,^46,47^ decompacts higher-order chromatin structures,^48,49^ and activates gene expression.^50,51^ Very recently, histone tail acetylation was shown to directly affect the viscoelastic properties of nucleosome condensates.^11^ Therefore, understanding the effects of individual histone post-translational modifications on chromatin condensability has strong implications for elucidating cellular functions.^29^

In parallel with experimental investigation, computational simulation is a powerful tool for quantitatively studying the dynamics and molecular details of chromatin organization. Numerous studies have been conducted across a wide range of resolution, from highly coarse-grained models^52–55^ to fully detailed atomistic simulations,^56–59^ including several studies on chromatin condensate formation.^6,10,22,55,60^ Nonetheless, detailed studies of the molecular origins of chromatin phase separation, condensate properties, and their modulation across different environments, particularly with respect to ionic strength and epigenetic modifications, remain limited. Understanding the impacts of these factors would significantly advance our knowledge of the genome architecture and its link to gene regulation. Residue-resolution coarse-grained models are uniquely suited for this purpose because they enable efficient simulations of large chromatin condensates while retaining the ability to investigate individual amino-acid-nucleotide interactions that contribute to phase-separating properties. ^61,62^

Here, we apply a residue-resolution chromatin model to study the phase-separating behavior of nucleosomal condensates. The model, implemented on a recently developed GPU platform,^63,64^ achieves high computational efficiency, enabling simulations of hundreds of nucleosomes over microsecond timescales within days while quantitatively reproducing multiple properties essential to chromatin interactions and organization. Our simulation reveals a non-monotonic dependence of nucleosome phase-separation propensity on salt concentration, with optimal condensation under physiological ionic conditions. Detailed analyses reveal that competition between global DNA-DNA repulsion and local histone-DNA attraction underpins this phase-separating behavior, which is further modulated by DNA conformational flexibility. Finally, we investigate the effect of histone tail acetylation and show that acetylation antagonizes nucleosome phase separation and accelerates nucleosome dynamics, establishing a direct contribution of epigenetic modifications to the regulation of chromatin organization.

## Results

### Residue-resolution model accurately captures chromatin properties

We have implemented a residue resolution chromatin simulation model on the GPU-accelerated OpenMM platform.^63,64^ The model combines a C*_α_*-based protein representation with a structure-based model^65,66^ for the ordered proteins and the molecular renormalization group (MRG) coarse-grained DNA model.^67^ Although the model does not include DNA sequence specificity, it quantitatively reproduces a series of properties essential for capturing chromatin organization. For example, it reproduces the experimentally reported persistence length of DNA at physiological salt concentration (Figure S1). The model also recapitulates inter-nucleosome interaction patterns and free energies obtained from an earlier, more detailed simulation model that has explicit ion representations^39^ (Figure S2). Most importantly, it reproduces experimentally measured sizes and shapes of higher-order chromatin structures under different conditions, as reflected by sedimentation coefficients across varying salt concentrations and linker lengths (Figure S3). All of these results demonstrate the model’s capability to capture ion-modulated chromatin structural transition.^68^ Additionally, this residue-level implicit-solvent model is highly efficient, simulating 100 nucleosomes for 1 *µs* (10^8^ timesteps) in 25 hours on a single L40S GPU, and can be further accelerated for multiple-replica simulations using the NVIDIA multi-process service (Figure S4). This is comparable to the state-of-the-art chromatin model at similar resolution.^60^

### Phase-separation of nucleosomes at physiological salt concentration

Since our model combines both residue-resolution accuracy for individual amino-acid-nucleotide interactions with high computational efficiency needed to simulate hundreds of nucleosomes at a reasonable cost, it allows us to systematically investigate nucleosome condensation and its modulation by various environmental factors. To investigate nucleosome condensation, we simulated 100 nucleosome core particles at physiological conditions (300K and 140 mM monovalent salt concentration). Consistent with experiments,^4,5,69^ nucleosomes phase-separate into condensates at physiological salt concentration (Figure 1), forming well-separated dilute and dense phases, with individual nucleosomes constantly exchanging between the two phases at the interface. Detailed analysis of nucleosomal conformations reveals striking differences depending on their local environment: nucleosomes in the dilute phase exhibit a fully wrapped structure, whereas those at the interface or within condensates exhibit partially unwrapped structures. Such an unwrapped breathing state of nucleosomes is important for histone-DNA interactions and may contribute to the stabilization of chromatin condensates.^6^

**Figure 1:**
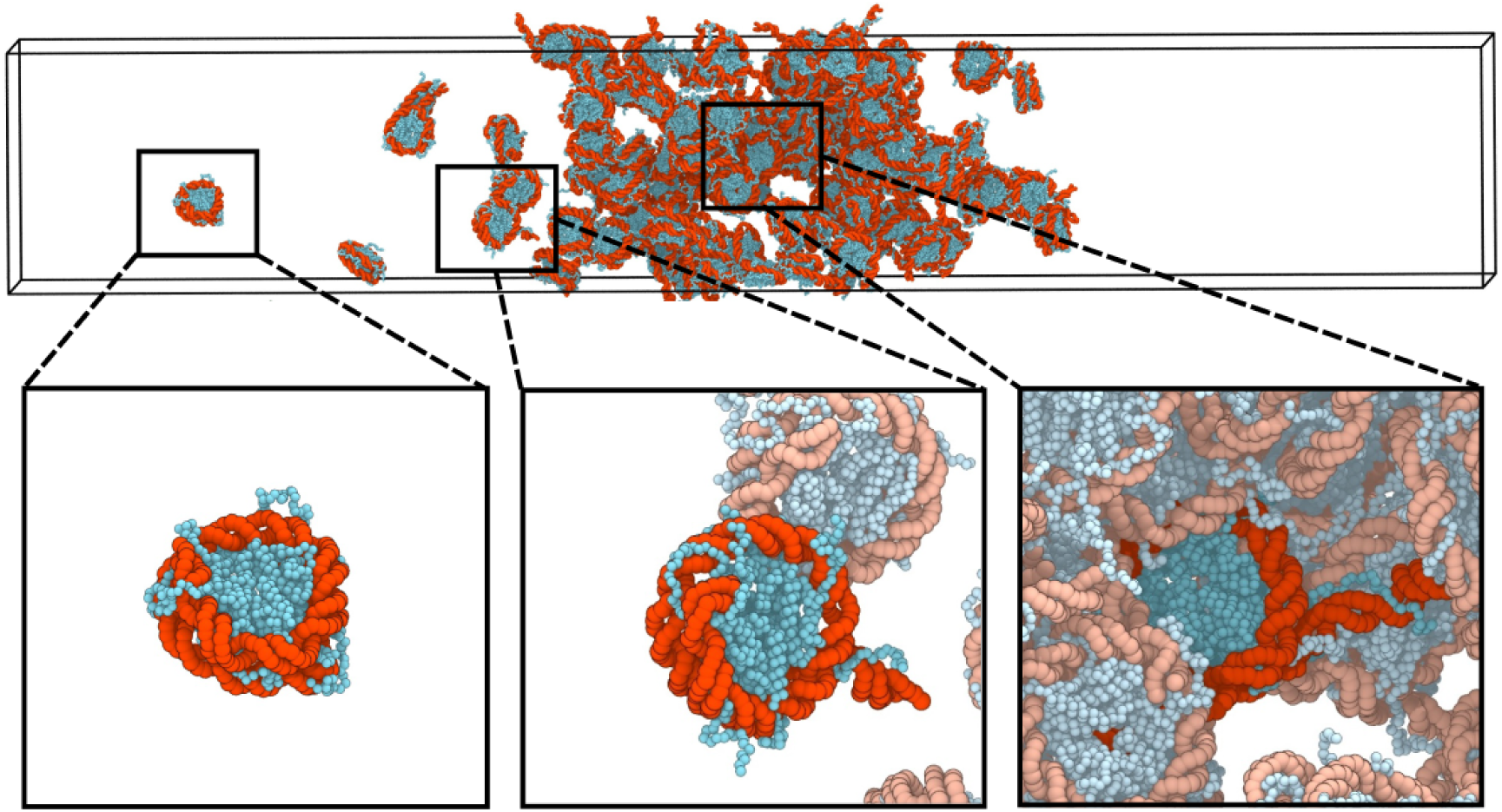
Multi-scale representation of the coarse-grained chromatin simulation model. Representative configuration from simulations of 100 nucleosome core particles within a slab geometry (49 × 49 × 300 nm^3^) at 140 mM monovalent salt concentration. The system is color-coded to distinguish its two primary components: DNA (red) and histone proteins (blue). Zoomed-in panels illustrate the three distinct conformational states observed across the system: “Outer” nucleosomes (located in the dilute phase) exhibit a fully-wrapped DNA configuration; “Inter” nucleosomes (located in the interfacial region) exhibit partial DNA unwrapping; and “Inner” nucleosomes (located in the condensate core) undergo extensive unwrapping, enabling a high density of internucleosomal interactions.

### Non-monotonic nucleosome phase separation is optimized at physiological salt concentration

Because the ionic environment is critical for chromatin organization and its regulation of downstream gene expression,^29,68,70,71^ we next examined nucleosome phase-separating behaviors across a range of salt concentrations. The simulations reveal a non-monotonic dependence of nucleosome phase separation on salt concentration, with optimal condensate formed under physiological conditions (Figure 2). The observation is consistent with an earlier turbidity experiment,^5^ which showed that nucleosome core particles phase-separate in buffer containing 150 mM NaCl, whereas condensates dissolve at both lower and higher salt concentrations.

**Figure 2:**
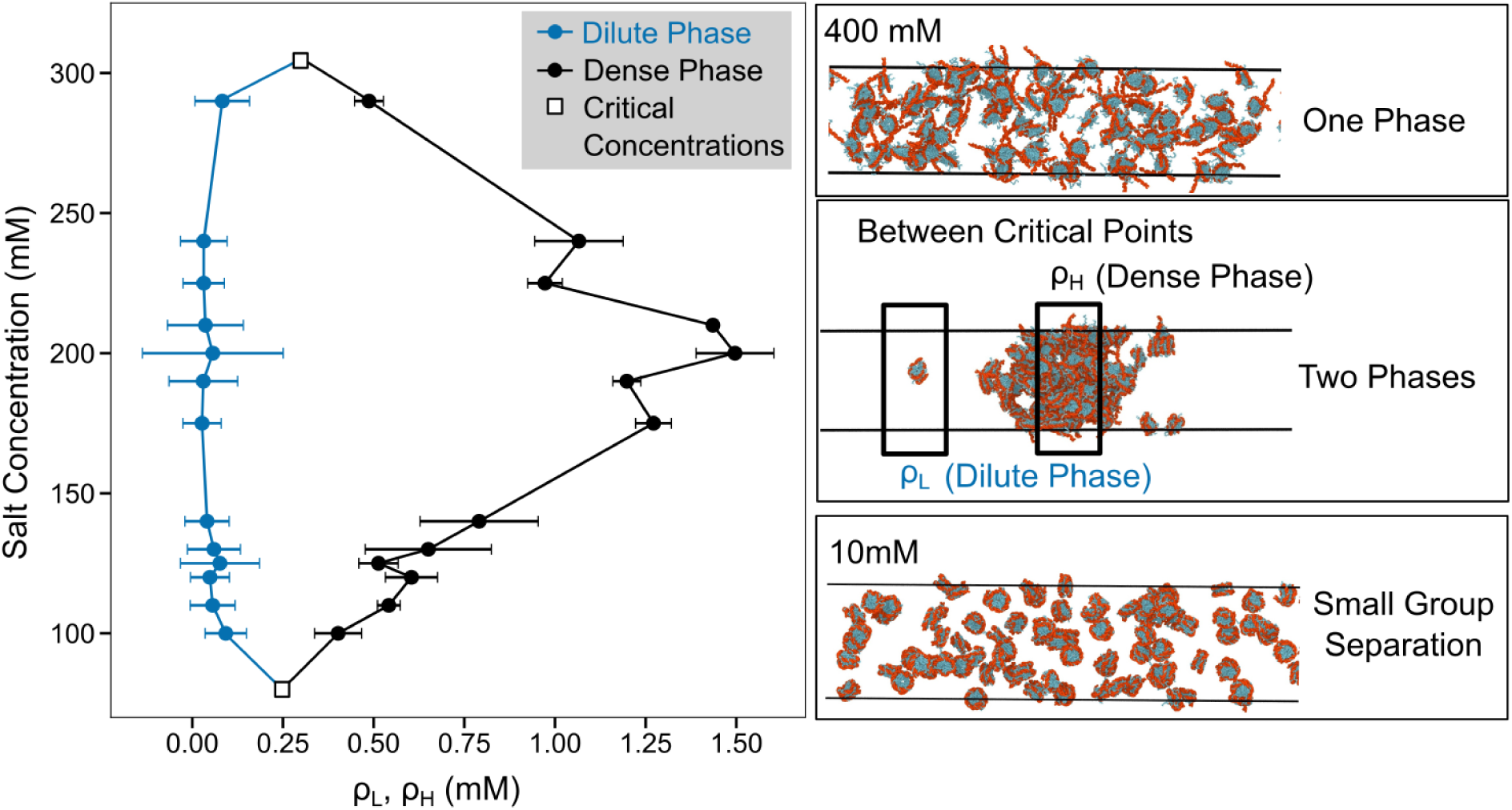
Phase behavior and salt-dependent morphological transitions of nucleosome condensates. (Left) Phase diagram of the nucleosome system. The binodal curve is defined by the salt-dependent densities of the dilute (blue line) and dense (black line) phases. The x-axis represents nucleosome density (mM), with the low-density regime (left) corresponding to the dilute phase and the high-density regime (right) representing the dense phase. Open squares denote the critical concentration points (*c*^∗^) marking the boundaries of the two-phase coexistence region. **(Right) Representative simulation snapshots** of the 100-nucleosome system across three distinct salt regimes. **Top (400 mM):** A single-phase state above the upper critical salt concentration (*c*^∗^_upper_), characterized by complete condensate dissociation. **Middle (175 mM):** A stable two-phase coexistence regime between the critical points, where the system partitions into a dense condensate core and a dilute surrounding phase. **Bottom (10 mM):** A microphase-separated regime observed below the lower critical salt concentration (*c*^∗^_lower_), where nucleosomes form small, kinetically-trapped oligomers rather than a bulk condensate.

Simulations across different salt concentrations allowed us to trace the coexistence of dense and dilute nucleosome phases and construct the corresponding phase diagram (Figure S5). Following earlier studies,^6,72,73^ we estimated the critical ionic concentrations based on the law of rectilinear diameter (Figures S6-S8, see Supplementary Section *Calculating Critical Concentrations* for details). This analysis identified an upper critical concentration of 300 mM salt and a lower critical concentration of 75 mM.

### Histone–DNA interactions drive nucleosome phase separation

Our fine-grained simulation model enables detailed examination of the residue-level interactions that drive the non-monotonic nucleosome phase separation propensity, particularly histone-DNA interactions (Figure 3A). To characterize these interactions, we computed protein–protein, protein–DNA, and DNA–DNA interaction energies among all nucleosomes in the simulation (Figure 3B). Consistent with the observed phase-separating behavior, attractive protein–DNA interactions exhibit a non-monotonic dependence on ionic concentration, reaching their maximum strength under physiological salt conditions. In contrast, protein–protein and DNA–DNA interactions are repulsive and vary relatively modestly over the same salt range. These results indicate that the sharp increase in protein–DNA interactions is the primary energetic driver of nucleosome phase separation. Thus, the simulated non-monotonic phase-separating behavior arises from the significant charge dichotomy, which polarizes the system through electrostatic interactions and drives a reentrant microphase-to-macrophase separation transition regulated by ion screening at distinct salt concentrations (See the *Discussion* section for more detail).

**Figure 3:**
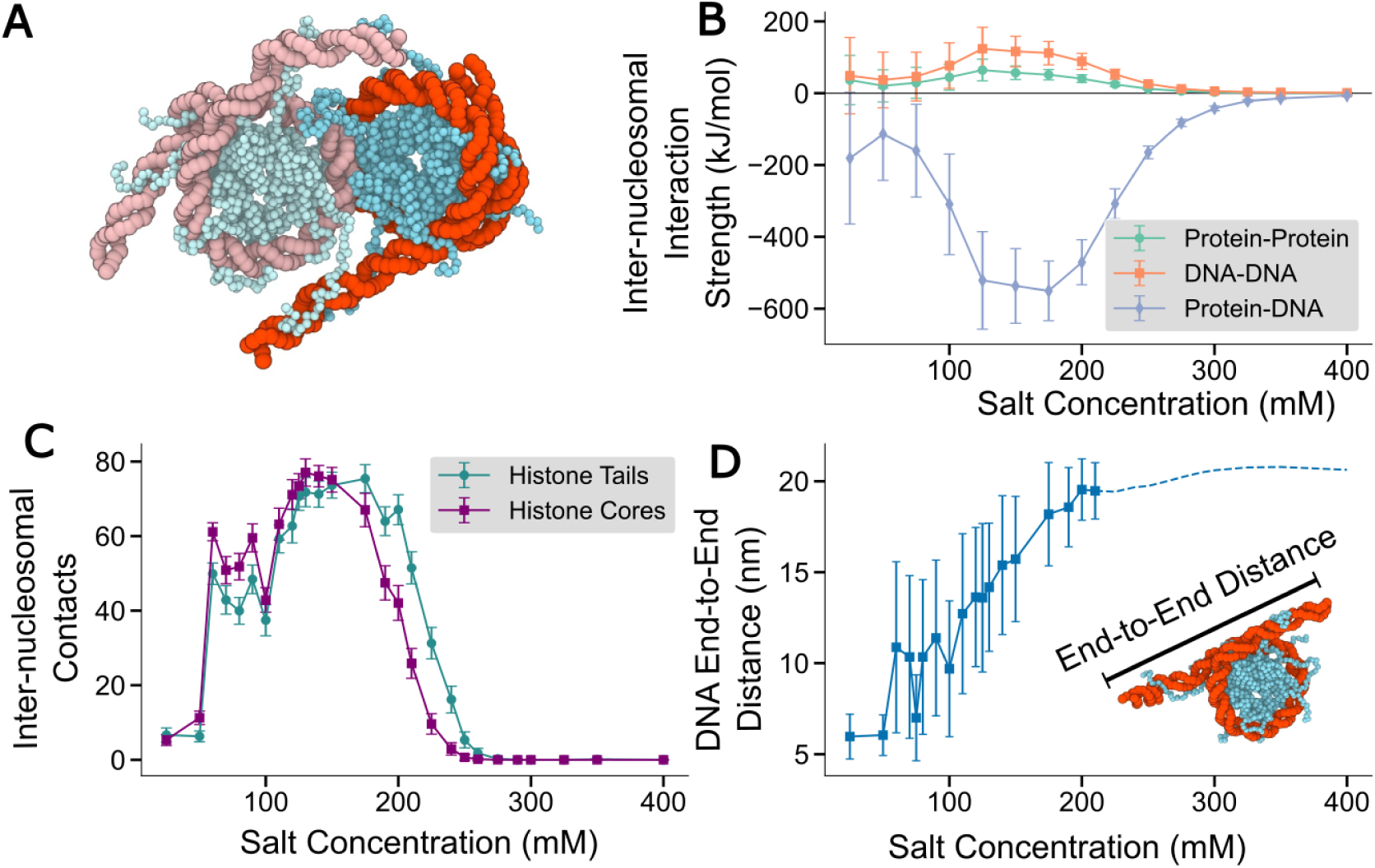
Energetic and structural drivers of inter-nucleosomal association. **(A) Schematic of inter-nucleosomal interactions.** Illustration of two interacting nucleosomes (primary nucleosome in standard colors; neighboring nucleosome shown in lighter colors for clarity). The highlighted region emphasizes the dynamic role of histone tails in bridging neighboring nucleosomes and promoting inter-nucleosome contact. **(B) Salt-dependent interaction energy profiles.** Average energetic contributions to inter-nucleosome interactions as a function of salt concentration. Repulsive DNA–DNA and protein–protein interactions (positive values) are outweighed by strong attractive protein-DNA interactions (negative values), which constitute the primary energetic driver for nucleosome condensation. **(C) Contact stoichiometry and valency.** Comparison of the number of inter-nucleosome contacts mediated by histone tail (teal)–DNA and histone core (purple)–DNA interactions across salt concentrations. The predominance of histone-mediated contacts underscores their central role as the primary “stickers” driving nucleosome condensate formation. **(D) DNA unwrapping dynamics.** Extent of Degree of DNA unwrapping (*nm*) as a function of salt concentration. The system achieves a maximum unwrapping of 20 nm, corresponding to the structural representation shown on the right.

To delineate the respective contributions from histone tails and histone cores to nucleosome phase separation, we analyzed the frequencies of inter-nucleosome contacts formed between these histone regions and nucleosomal DNA (see Method Section *Computational of Inter-nucleosome Contacts* for details). Interestingly, histone core–DNA interactions follow the same salt-dependent trend as histone tail–DNA interactions, but make a larger energetic contribution at lower salt concentration (Figure 3C), indicating that histone cores also contribute substantially to phase separation. This finding is consistent with simulations performed using our explicit-ion residue-level model^39^ and with earlier chromatin phase-separation simulations using a nucleosome-level coarse-grained model,^55^ showing nucleosome face–side interactions significantly contribute to inter-nucleosome association and chromatin phase separation.

Because chromatin DNA contributes significantly to chromatin phase separation,^4,6^ and because our simulations reveal significant differences in nucleosomal DNA wrapping between the dilute- and dense-phase nucleosomes (Figure 1), we hypothesize that nucleosomal DNA unwrapping strengthens protein-DNA interactions under physiological conditions. To assess the contribution of nucleosomal DNA unwrapping to nucleosome phase separation, we computed the average end-to-end distance of nucleosomal DNA across salt concentrations (Figure 3D). As expected, DNA unwrapping increases with salt concentration, leading to stronger protein–DNA interactions that peak at physiological ionic strength and correlate with the propensity of nucleosomes to phase separate. At higher salt concentrations, ion screening becomes dominant, and all interactions gradually weaken toward neutrality. Since the inner wrap of nucleosomal DNA was regidified in our simulations to improve computational efficiency (See Methods Section *Details of Residue-resolution Chromatin Model* for details), we could not evaluate the contribution of inner-layer DNA to condensate formation. We expect that releasing this restraint would further favor condensate formation at higher salt concentrations.

### Salt-dependent Cluster Formation

The conformations of charge-imbalanced polyampholytes are governed by a delicate competition between short-range Coulombic attractions and long-range repulsions, leading to distinct structural regimes dependent on charge asymmetry and ionic strength.^40^ While our simulations use disconnected nucleosome monomers, each nucleosome functions as a modular polyelectrolyte-polyampholyte hybrid, in which disordered histone tails act as polyampholytic brushes that interact with the high-charge-density DNA scaffold. This hybrid architecture suggests that the inter-nucleosomal interactions driving clustering are subject to the same polarization-induced attractions and surface-charge-patch effects observed in other electrostatic-driven complex formation.^41^

In cells, nucleosomes form discrete, heterogeneous assemblies known as nucleosome clutches. ^74^ The size and density of these clutches are cell-type specific and correlate with functional states of chromatin; for instance, smaller, less dense clutches are characteristic of pluripotent stem cells, whereas larger and denser clutches are stabilized by extrinsic factors such as linker histone H1.^74–76^ To examine how ionic strength regulates cluster organization, we analyzed phase-separated nucleosomes across a range of salt concentrations (Figure 4). Our analysis reveals a non-monotonic trend: nucleosomes form small clusters at low salt, coalesce into a single large cluster at physiological salt, and dissolve again at high salt. This behavior mirrors the reentrant phase transitions predicted for polyampholytes, where salt initially screens repulsions to promote aggregation but eventually screens the attractions required for cluster stability.^40,41^ Within these clusters, nucleosomes formed interlined networks via histone–DNA interactions, thereby stabilizing the condensate (Figure S9, see Methods Section *Computational of Inter-nucleosome Contacts* for details). Graph-theoretic analysis indicates that these clusters adopt a “small-world” network topology, characterized by a percolated architecture of physical crosslinks^77^ (Figure S10, see Methods Section *Computation of Nucleosome Networks* for details). Such an organization suggests that the nucleosome condensate forms inhomogeneous networks where connectivity and “hub” nucleosomes play critical roles in defining the condensate rheology and internal stability.^77^

**Figure 4:**
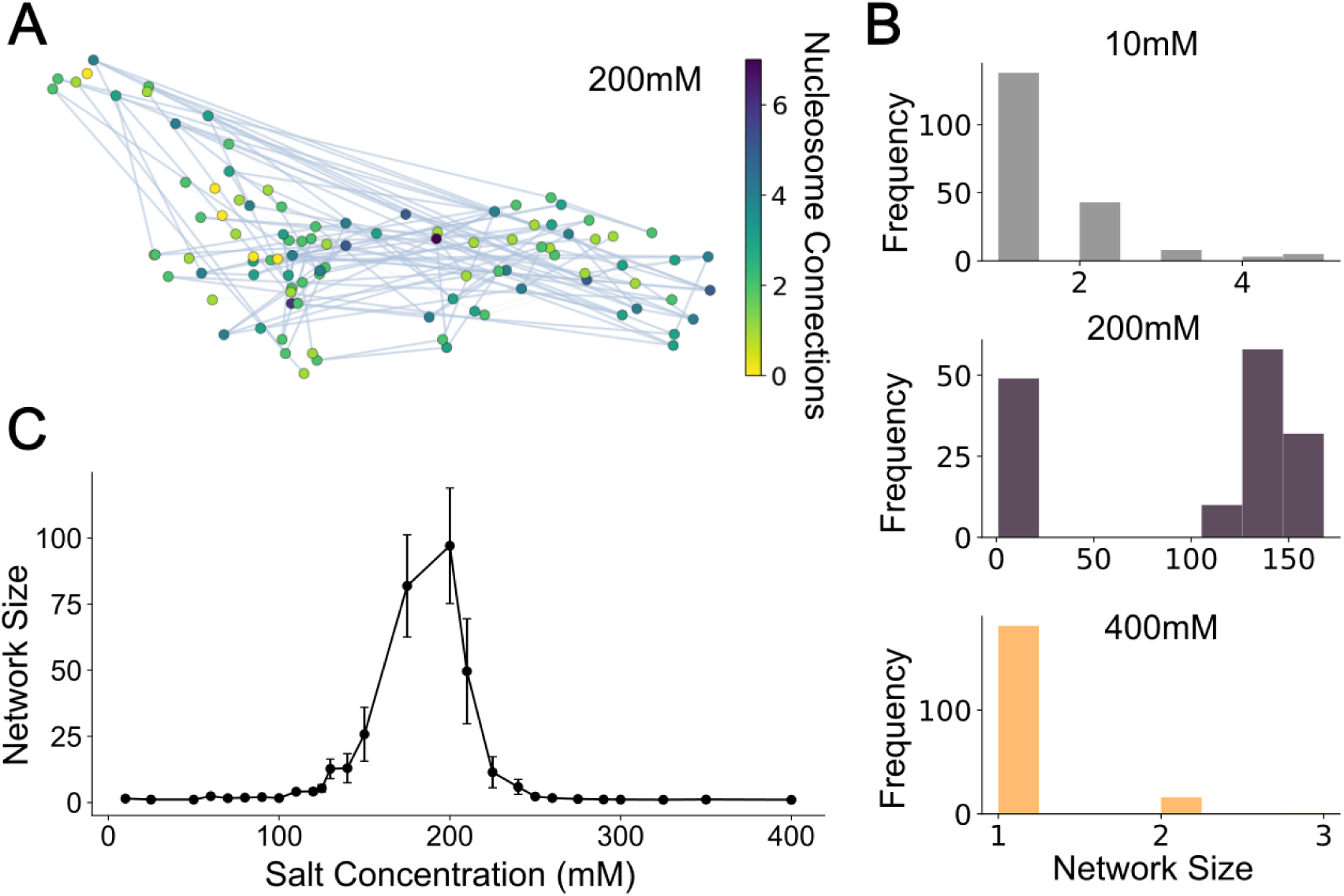
Topological connectivity and network scaling within the chromatin condensate. **(A) Representative nucleosome interaction network at 200 mM.** Representative simulation snapshot where nodes correspond to nucleosomes and edges denote inter-nucleosomal contacts. Nodes are colored by degree of connectivity using the Viridis scale: yellow nodes represent isolated nucleosomes (degree 0), whereas darker purple nodes represent highly connected “hubs” within the condensate core. **(B) Distribution of network cluster sizes across salt regimes.** Stacked histograms showing the frequency of cluster sizes, defined as the total number of connections per cluster, over the full simulation trajectory. At 10 mM (grey), nucleosomes undergo small-scale oligomerization, with clusters restricted to 1—6 connections. At 200 mM (purple), the system forms a large-scale interaction network with a broad distribution reaching *>* 150 connections, indicative of a highly interconnected macroscopic condensate. At 400 mM (yellow), the system returns to a predominantly monomeric state, with clusters reaching at most 2 connections, reflecting the absence of stable inter-nucleosomal contacts. **(C) Average network connectivity as a function of salt concentration.** Mean cluster size calculated across all simulation snapshots for each salt concentration. Connectivity reaches maximum at 200 mM, identifying the optimal salt regime for multivalent network percolation.

### Influence of DNA Conformational Flexibility on Nucleosome Phase Separation

The conformational flexibility of unwrapped DNA may facilitate chromatin phase separation by creating a heterogeneous environment that excludes intruding molecules. ^78^ To investigate the influence of DNA flexibility on nucleosome phase separation, we performed simulations with varying DNA rigidity by adjusting the strength of bonded interactions (See Method Section *Tuning DNA flexibility in Simulation* for details). We found that DNA mechanical properties significantly alter nucleosome phase-separating behavior: increasing DNA rigidity promotes nucleosome condensate formation and increases nucleosomal densities in the dense phase (Figures 5A and B). Consistent with our previous analyses (Figure 3), more rigid DNA unwraps to a greater extent from nucleosomes (Figure S11), leading to stronger interactions with histone proteins that drive condensate formation (Figures 5C and D). Because DNA rigidity depends on sequence composition,^79–82^ these results suggest a potential mechanism by which DNA sequence specificity can modulate nucleosome condensation and, ultimately, genome organization.

**Figure 5:**
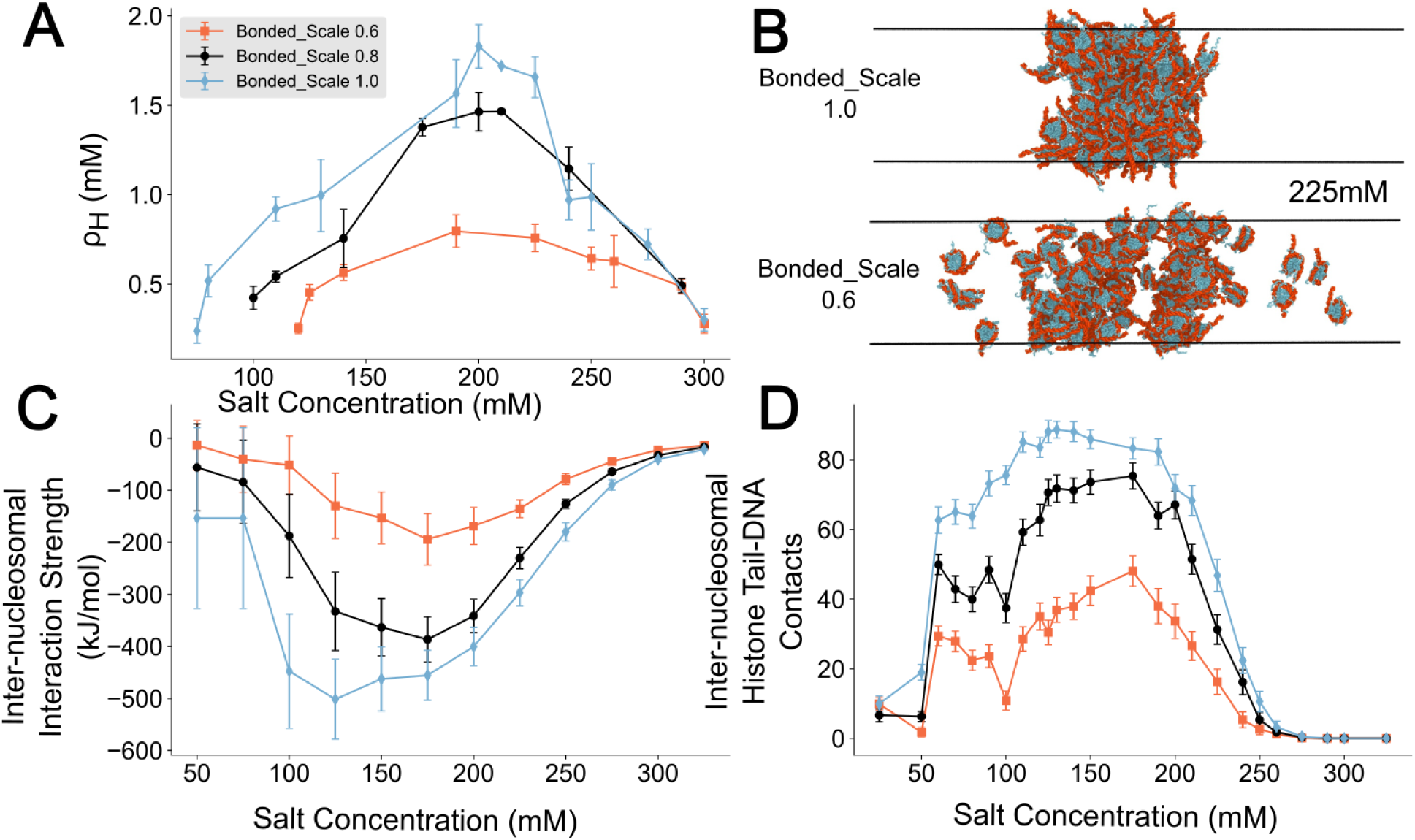
DNA bending rigidity modulates condensate density and interaction stoichiometry. **(A) Salt-dependent density of the condensed phase.** Dense-phase density (*ρ_H_*) is plotted as a function of salt concentration for three levels of DNA bending rigidity: Bonded scale = 0.6 (orange), 0.8 (unmodified, black), and 1.0 (light blue). Increasing DNA rigidity enhances phase-separating propensity and significantly increases the maximum condensate density across salt concentrations. **(B) Morphological sensitivity to polymer mechanics.** Representative simulation snapshots at 225 mM NaCl comparing high-rigidity (Bonded scale = 1.0, top) and low-rigidity (Bonded scale = 0.6, bottom) regimes. The arrow indicates increasing DNA rigidity, which is associated with greater condensate compaction and a more well-defined phase interface. **(C) Global interaction energetics.** Total interaction energy, calculated as the sum of protein–protein, DNA–DNA, and protein–DNA energetic contributions. Rigid DNA (Bonded scale = 1.0) strengthens attractive interactions within the system, whereas flexible DNA (Bonded scale = 0.6) weakens stabilizing interactions, resulting in a less cohesive condensate. **(D) Tail-mediated contact valency.** Number of inter-nucleosomal histone tail–DNA contacts across salt concentrations. Higher DNA rigidity increases contact frequency, supporting a model in which stiffer DNA facilitates exposure and accessibility of histone “stickers” to their binding partners.

### Acetylation-modulated nucleosome phase separation

Acetylation of lysine residues on histone tails is a key histone post-translational modification associated with gene activation.^50^ Acetylation can disrupt chromatin stability,^45,83^ weaken inter-nucleosomal interactions,^47^ and decompact higher-order chromatin organization.^48,49^ Recent studies have shown that histone tail acetylation, particularly on the H3 and H4 tails, reduces the propensity of chromatin to undergo phase separation.^4,11,69^ To investigate how acetylation of key histone tails affects nucleosome phase separation, we modeled acetylated H3 and H4 tails by neutralizing the corresponding positively charged lysine residues. Consistent with experimental observations, our simulations show that acetylation of either H3 or H4 tails dissolves the phase-separated nucleosome condensate (Figure 6A), leading to a reduced dense-phase density relative to wild-type nucleosomes (Figures 6B and S12). Different from previous experiments reporting a strong effect of H4 tail acetylation on chromatin phase separation,^69^ our simulation shows that H3 tail acetylation has a more pronounced effect in dissolving nucleosome condensates than H4 tail acetylation. The difference may be attributed to the absence of linker DNA in our simulations. Although H3 histone tails are generally thought to mediate intra-nucleosomal interactions,^84,85^ the absence of linker DNA and neighboring connected nucleosomes in our nucleosome core particle simulations likely promotes H3-tail interactions with other nucleosomes, thereby contributing significantly to condensate formation. Consistent with this interpretation, H3 tail acetylation causes a substantial reduction in inter-nucleosomal contacting frequency (Figure 6C).

**Figure 6:**
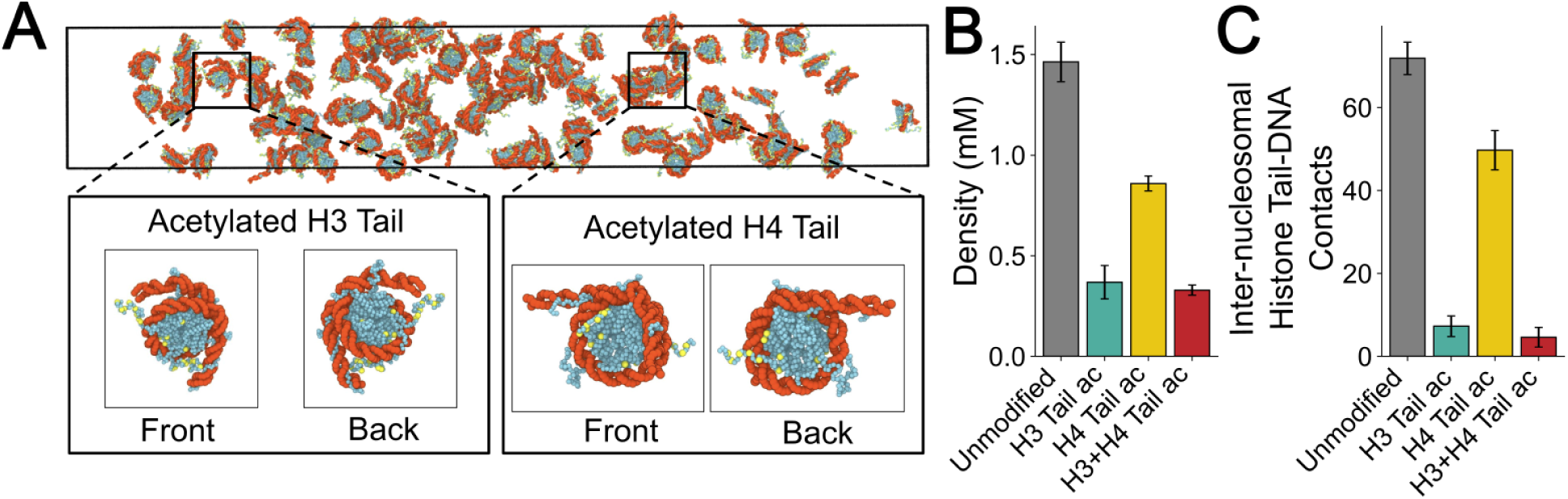
Histone tail acetylation modulates condensate density and multivalent connectivity. **(A) Structural representation of acetylated nucleosomes.** Representative simulation snapshot at 200mM NaCl. Neutralized lysine residues in the histone tails are highlighted in yellow to represent the acetylated state, while DNA (red) and the histone octamer (blue) follow the established color scheme. Insets: Magnified views of the H3 (left) and H4 (right) histone tails, illustrating the spatial distribution of acetylation sites on opposite sides of the nucleosome core. **(B) Effect of tail-specific acetylation on condensate density.** Dense-phase density (*ρ_H_*) at 200mM salt is compared across four conditions: unmodified (grey), H3 tail acetylation (teal), H4 tail acetylation (yellow), and combined H3 and H4 tail acetylation (red). Acetylation progressively reduces condensate compaction, with combined H3/H4 acetylation producing the most dilute dense phase. **(C) Stoichiometry of inter-nucleosomal tail–DNA interactions.** Total number of inter-nucleosomal histone tail-DNA contacts at 200mM salt. The trend mirrors the density profiles in panel B, where the unmodified system (grey) exhibits the highest connectivity, followed by H4 tail acetylation (yellow), H3 tail acetylation (teal), and combined H3 and H4 tail acetylation (red). These results indicate that charge neutralization reduces histone-tail “sticker” valency, thereby weakening the global connectivity of the chromatin network.

### Dynamic nucleosomes within nucleosome condensates

The importance of H3 tails in regulating nucleosome condensate formation was recently demonstrated by a comprehensive microrheology study of different H3 tail mutants,^11^ which showed a significant increase in nucleosome viscosity upon mutation of H3 basic residues. A similar trend was also observed in coarse-grained simulations of different H3 tails with DNA.^11^ Motivated by these findings, we computed the viscosity of nucleosomes in our condensate simulation at a salt concentration of 200 mM (Figure S13), comparing unmodified nucleosomes with nucleosomes bearing acetylated H3 and H4 tails. Because the timescale of residue-resolution coarse-grained models is not rigorously defined due to the use of a small friction constant (*γ* = 0.01 *ps*^−1^, approximately 100-fold smaller than water viscosity),^86^ the absolute values of the computed diffusion coefficient should be interpreted with caution. Nonetheless, our simulations show that acetylation of H3 and H4 tails significantly reduces nucleosomal viscosity within condensates while increasing diffusion (Figure 7), consistent with the trends reported in the microrheology experiment.^11^

**Figure 7:**
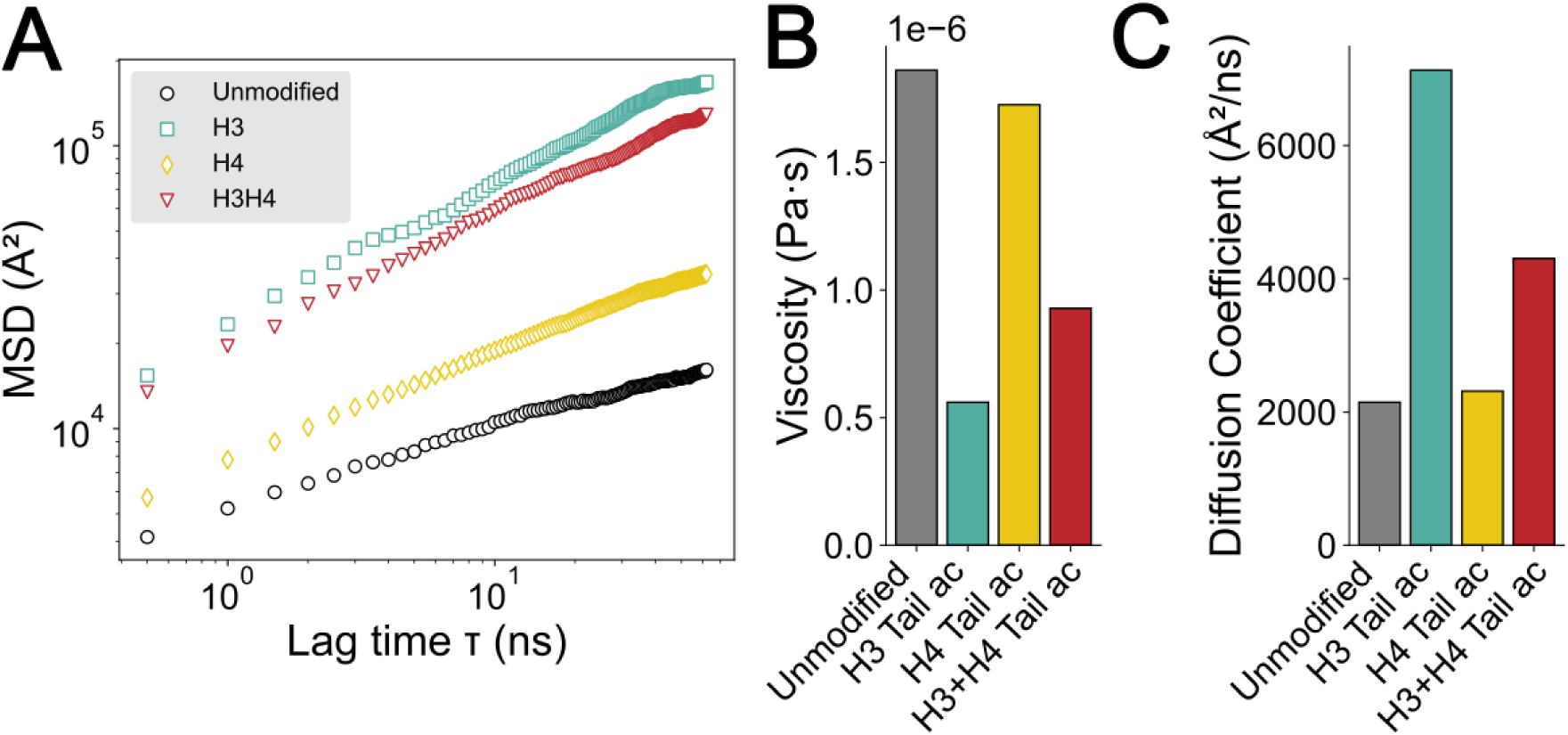
Dynamic and rheological properties of acetylated nucleosome condensates. **(A) Mean Squared Displacement (MSD) of nucleosomes.** MSD as a function of time for unmodified (grey), H3 tail-acetylated nucleosomes (teal), H4 tail-acetylated nucleosomes (yellow), and nucleosomes with both H3 and H4 tail acetylation (red) at 200 mM physiological salt concentration. The slopes of the MSD curves reveal a transition from a sub-diffusive, “glassy” regime in the unmodified condensates to a more fluid, mobile state upon tail acetylation. **(B) Condensate viscosity (***η*). Nucleosomal viscosity compared across the four acetylation states. Acetylation, particularly combined H3 and H4 tail acetylation, significantly decreases condensate viscosity, indicating that weakened histone tail–DNA interactions (Figure 6) promote a more liquid-like and less mechanically resistant material state. **(C) Diffusion coefficients (***D*). Calculated diffusion coefficients of nucleosomes within the dense phase. The increased mobility observed upon combined H3 and H4 tail acetylation (red), relative to the unmodified system (grey), demonstrates that epigenetic modifications can regulate the internal dynamics of nucleosome condensates, potentially modulating transcriptional activity.

## Discussion

The nucleosome is a polyampholyte, with a significant population of positive charges on the histone proteins (+148*e*) and negative charges on the DNA (−294*e*), resulting in a net negative charge of −146*e* per nucleosome. The charge imbalance leads to two competing electrostatic effects: global repulsion between negatively charged nucleosomes and local attraction between histone proteins and DNA. The non-monotonic dependence of nucleosome condensation on salt concentration can therefore be understood as the outcome of competition between these two types of interactions. At intermediate salt concentrations, DNA-DNA repulsions are sufficiently screened, while attractive histone-DNA attractions remain within the corresponding electrostatic screening length, leading to phase separation.^41^ At lower salt concentrations, both attractive and repulsive interactions remain long-ranged, and the dominant repulsion between nucleosomes prevents the formation of large condensates. Nonetheless, localized microphase separation can still occur,^40^ driven by strong localized histone-DNA attractions. At high salt concentrations, both attractive and repulsive electrostatic interactions are strongly screened, leading to dissolution of the condensates into individual nucleosomes. The strong screening of charge effects is further reflected in the simulated nucleosome structures (Figures 2 and 3D), where high-salt conditions induce substantial DNA unwrapping. Notably, the residue-resolution of our model is well-suited for resolving this concentration-dependent phase-separating behavior, because its effective interactions fall within the length scales of the Debye-Hűckel length at corresponding salt concentrations.^41,87^

The onset of large condensates is often preceded by the formation of subsaturated clusters, forming a percolation network and macroscopic phase separation.^88–90^ Our clustering analysis supports this mechanism, showing that the largest nucleosome contacting networks form under physiological salt conditions (Figure 4). Interestingly, at low salt concentrations, such as 10mM, nucleosomes do not assemble into large condensates but instead become enriched in highly segregated small clusters that resemble microphase separation in polymeric materials.^87,91–93^

Our simulations also demonstrate the importance of DNA flexibility in nucleosome phase separation. As DNA rigidity increases, nucleosomes show a greater propensity to form condensates. This enhanced condensation may come from three related effects. First, the transition from the dilute phase to the dense phase incurs an entropic cost associated with restricting DNA conformational mobility. Increased DNA rigidity reduces DNA conformational entropy in the dilute state, thereby lowering the entropic penalty of DNA incorporation into the dense phase and favoring condensate formation. Second, more rigid DNA unwraps to a greater extent (Figure S11), exposing more histone protein surface and increasing multivalent interactions with DNA (Figure 5C), thereby promoting condensation. Third, increasing DNA rigidity increases the effective correlation length of a nucleosome,^78^ which can reduce the free energy penalty for inserting additional nucleosomes into an existing cluster, thereby promoting the growth and stability of the nucleosome condensate.

Our efficient GPU-implemented residue-resolution model quantitatively reproduces several chromatin structural and interaction properties and achieves approximately 1000 steps per second (Figure S4), corresponding to ∼ 1*µs* per day on a single GPU for a 100-nucleosome system. This computational efficiency enables studies of large chromatin systems involved in phase separation and genome organization.^4,88,94^ At the same time, the residue-level resolution of the model allows detailed examination of interactions modulated by post-translational modifications at individual residues. We have shown that histone tail acetylation reduces the propensity of nucleosomes to phase separate and increases their dynamics (Figures 6 and 7). One future direction is to incorporate protein-DNA sequence specificity into the model framework,^82,95–99^ because heterotypic interactions are essential to drive biomolecular phase separation.^100–102^ Additionally, combined with other residue-resolution models,^72,97,103–106^ this model will enable mechanistic studies of other biomolecules important for chromatin and epigenetic regulation.^42,44^

## Methods and Materials

### Details of Residue-resolution Chromatin Model

We employed a residue-resolution coarse-grained chromatin model with implicit treatment of water and ions. Detailed descriptions of this model can be found in previous publications.^73,107^ The DNA was modeled with a coarse-grained Molecular Renormalization Group (MRG-CG) model.^67,108^ Proteins were modeled using the maximum entropy optimized force field (MOFF) model.^103,109^ This combined DNA-protein model has previously been applied to simulations of several protein-DNA LLPS systems, including heterochromatin protein 1 (HP1)-DNA^73^ and linker histone H1-DNA systems,^107^ successfully reproducing experimentally measured phase-separation propensities. In this manuscript, we apply this model, implemented on the GPU-accelerated OpenMM platform,^64,76^ to chromatin and validate it using higher-order chromatin organization and phase-separation behavior.

The MOFF force field has been documented in detail in previous publications.^63,73,103,107,109^ Briefly, proteins are modeled at residue resolution, where each amino acid is represented by a single bead at the *α*-carbon position. The total potential energy governing protein-protein interactions is defined as:

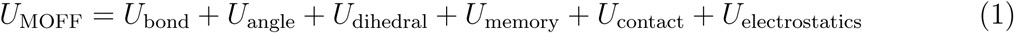

where *U*_bond_, *U*_angle_, and *U*_dihedral_ maintain the protein backbone geometry, using the following forms:

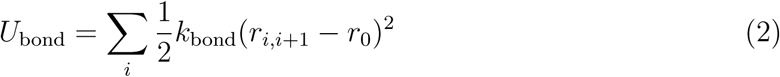

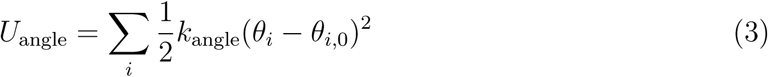

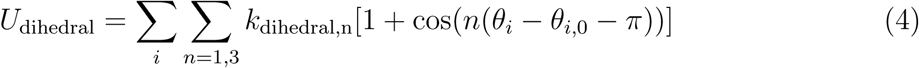

where *r*_0_ = 0.38 nm, *k*_bond_ = 1000 kJ/mol/nm^2^, *k*_angle_ = 120 kJ/mol/rad^2^, *k*_dihedral,1_ = 3.0 kJ/mol/rad^2^, and *k*_dihedral,3_ = 1.5 kJ/mol/rad^2^.

The memory term, *U*_memory_, is a native-pair potential that stabilizes the ordered secondary and tertiary structures of folded protein domains based on the input configurations.

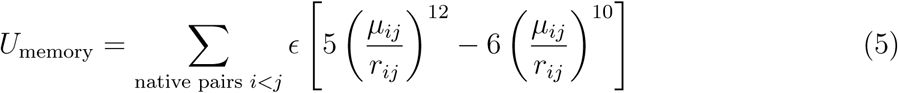

This term is applied only to native pairs identified from the initial input native structure using the Shadow Algorithm,^110^ where *µ_ij_* is the distance between *C_α_* atoms from residues *i* and *j*, and *ɛ* = 3 kJ/mol. Non-bonded residue-specific interactions are governed by a pairwise contact potential, *U*_contact_, derived via an iterative maximum entropy optimization algorithm to achieve consistent accuracy for both ordered and disordered proteins.^103^ The contact potential is given by

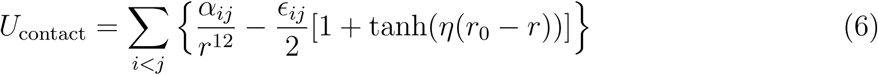

where the repulsive 1*/r*^12^ term describes the excluded-volume interactions between simulated particles, and the tanh-based term describes residue-specific protein-protein interactions optimized via maximum entropy optimization.

Electrostatic interactions, *U*_electrostatics_, are modeled using a Debye-Hűckel potential with a functional form:

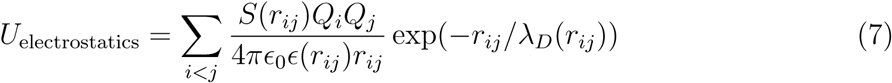

which incorporates a distance-dependent dielectric constant, *ɛ*(*r*), to capture changes in local solvation environment:

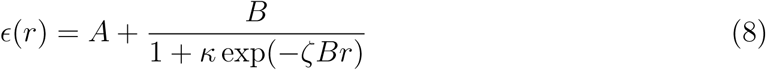

This potential gradually decays to zero at a cutoff distance of at least three times the corresponding Debye-Hűckel screening length.

To improve computational efficiency, the histone core residues and the inner wrap of nucleosomal DNA were rigidified during the simulation. Our previous studies^39,76,111^ have demonstrated that rigidified nucleosome models quantitatively reproduce chromatin properties obtained from simulations of unrigidified nucleosomes.

The MRG force field has also been documented in detail in previous publications.^67,108^ Briefly, each nucleotide is represented by a single coarse-grained site with a mass of 325 Da. The DNA energy function is defined as:

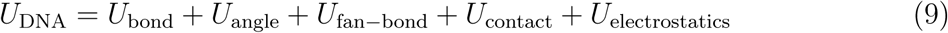

The bonded potentials among coarse-grained sites within a single DNA strand are defined by *U*_bond_ and *U*_angle_, with the functional form:

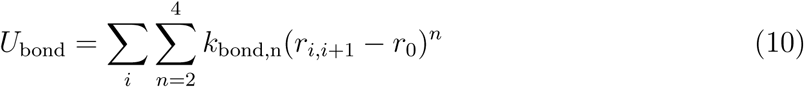

and

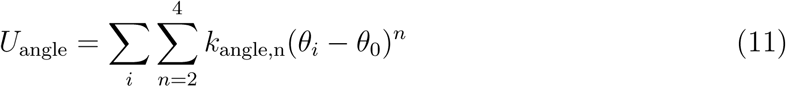

The fan bonds potential, *U*_fan−bond_, was defined between two ssDNA strands, using equation:

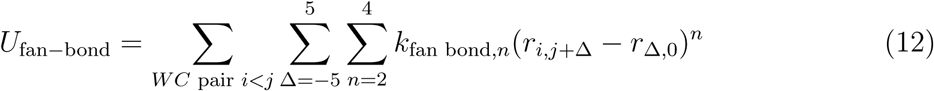

This term captures base-pairing and cross-stacking interactions between the two strands. To accurately model the mechanical properties of DNA, bonded interactions were scaled by a factor of 0.8 to reproduce experimental persistence lengths (Figure S1). The electrostatic potential was updated to the same distance-dependent Debye-Hűckel form as in MOFF, with each nucleotide carrying a charge of −1*e*.

The corresponding parameters are listed below:

Protein-DNA interactions include both contact and electrostatic potentials. The contact potential represents excluded-volume interactions and is defined as

**Table 1:** Parameters for DNA bonded potential. All the *k_bond,n_* are reported in kcal/mol/nm^2^, and *r*_0_ is reported in nm.

| $k_{\text{bond},2}$ | $k_{\text{bond},3}$ | $k_{\text{bond},4}$ | $r_0$ |
| --- | --- | --- | --- |
| 262.5 | -226 | 149 | 0.496 |

**Table 2:** Parameters for the DNA angle potential: All *k*_angle,n_ are reported in kcal/mol/degree^2^, and *θ*_0_ is reported is degree.

| $k_{\text{angle},2}$ | $k_{\text{angle},3}$ | $k_{\text{angle},4}$ | $\theta_0$ |
| --- | --- | --- | --- |
| 9.22 | 4.16 | 1.078 | 156 |

**Table 3:** Parameters for the DNA fan-bond potential. All *k*_fan-bond,n_ are reported in kcal/mol/nm^2^, and *r*_Δ,0_ is reported in nm. Here, Δ defines the fan bond between coarse-grained nucleotide *i* and *j* + Δ, where nucleotide *i* and *j* are Watson–Crick (WC)-paired

| $\Delta$ | $k_{\text{bond},2}$ | $k_{\text{bond},3}$ | $k_{\text{bond},4}$ | $r_{\Delta,0}$ |
| --- | --- | --- | --- | --- |
| -5 | 4.67 | 2.1 | 1.46 | 1.71 |
| -4 | 0.0001324 | -12.2 | 18.5 | 1.635 |
| -3 | 8.5 | -44.4 | 50.0 | 1.47 |
| -2 | 12.3 | -40.0 | 37.0 | 1.345 |
| -1 | 4.0 | -10.0 | 8.0 | 1.23 |
| 0 | 292.0 | 410.0 | 720.0 | 1.13 |
| 1 | 11.5 | -41.0 | 58.0 | 0.99 |
| 2 | 9.55 | -45.9 | 50.2 | 0.92 |
| 3 | 13.78 | -52.7 | 50.0 | 1.02 |
| 4 | 13.86 | -56.8 | 50.0 | 1.25 |
| 5 | 36.26 | -77.0 | 50.0 | 1.69 |

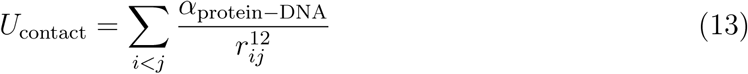

where *i* and *j* denote coarse-grained protein and DNA beads, respectively, and *α*_protein−DNA_ = 1.6264 × 10^−3^ nm^12^ · kJ/mol. Electrostatic interactions are modeled using the same distance-dependent Debye-Hűckel potential used for protein-protein and DNA-DNA interactions.

### Details of Simulation Setup

#### Simulation System Setup

The simulation systems, including single-nucleosome, two-nucleosome, and 12-mer chromatin, were modeled using the same protocol as described in our previous publications.^39,76^ Initial configurations were constructed based on the crystal structures of single nucleosomes (PDB ID: 1KX5,^112^ 3LZ1^113^) and tetranucleosomes,^114^ and used the 3DNA software package^115^ to build and align linker DNA to connect adjacent nucleosomes. In addition, we constructed a 12-mer nucleosome array with a nucleosome repeat length of 177 base pairs (Figure S3).

#### Simulation Details

All simulations were carried out in an implicit-solvent environment using the OpenABC software package,^63^ implemented on the OpenMM platform.^64^ Simulations were performed at 300 K using the Langevin Middle Integrator^116^ with a friction coefficient of 0.01 ps^−1^ and a time step of 10 fs. To improve computational efficiency, the inner 73-bp layer of nucleosomal DNA and the histone core were rigidified throughout the simulations, while the outer 74-bp layer of nucleosomal DNA remained flexible to preserve the plasticity needed for capturing nucleosome breathing and DNA wrapping dynamics.^6^

#### Phase Separation Simulation

Phase-separation simulations were performed using systems consisting of 100 single nucleosomes in a slab geometry of 50 × 50 × 300 nm^3^) with periodic boundary conditions. To prepare the phase-separation simulation, nucleosomes were first equilibrated in a cubic box (100 × 100 × 100 nm^3^) for 420 ns. The system was then compressed under an isothermal-isobaric (NPT) ensemble at 150 K and 1 bar to promote nucleosome condensation. The compressed box was subsequently elongated along the *z* direction to reach 300 nm. Production runs were performed under a canonical (NVT) ensemble for 1.05 *µ*s across different salt concentrations. The final 250 ns of each trajectory was used to analyze nucleosome clustering, network topology, and transport properties.

#### Two-nucleosome Simulations

Following previous studies,^39,49^ simulations of inter-nucleosome interactions were carried out using two nucleosome core particles with 601-sequence DNA, without linker DNA or linker histone. To enhance the sampling of the binding free energy landscape, we used umbrella sampling^117^ implemented via the PLUMED software package.^118^ Two collective variables (CVs) were used, the first, *r*, is the distance between the geometric centers of two nucleosomes, defined using residue IDs: 63-120, 165-217, 263-324, 398-462, 550-607, 652-704, 750-811, and 885-949 from the nucleosome crystal structure (PDB ID: 1KX5).^119^ The second CV, *θ*, was defined as the angle between two unit vectors perpendicular to the nucleosome faces. The umbrella restraints were applied with a spring constant of 5.0 kJ/(mol·nm^2^) for *r* and 0.005 kJ*/*(mol · ◦^2^) for *θ*. Umbrella centers were placed on a uniform grid, with *r* ranging from 6.0 to 13.0 nm, and *θ* from 0^◦^ to 180^◦^, using intervals of 1.0 nm and 30^◦^. Each umbrella trajectory was simulated for 24 million steps with a 10 fs time step. We excluded the first 12.5 million steps of each trajectory when constructing the unbiased binding free energy profile (Figure S2A, black).

For comparison, the same two-nucleosome umbrella simulation protocol was also performed using a more detailed explicit-ion model^39^ at 150 mM NaCl. Due to significantly greater computational complexity, we performed 1.3 million steps per umbrella window and excluded the first 3 million steps when reconstructing the unbiased binding free-energy profile (Figure S2A, red).

#### 12-mer Chromatin Simulation

To fully sample the conformation space of the 12-mer nucleosome array, we performed umbrella sampling^117^ using the collective variable (CV) *Q*, computed with PLUMED.^118^ The CV *Q* quantifies the similarity between a simulated chromatin conformation and a reference two-start chromatin structure^120^ and is defined as:

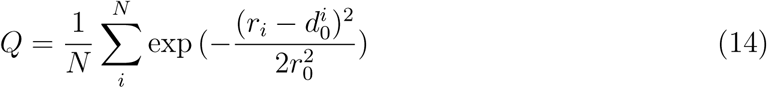

In this equation, *N* = 66 is the total number of nucleosome pairs in the 12-mer array, *r_i_* is the distance between nucleosomes in the *i*th pair, *d^i^*_0_ is the corresponding distance in the reference two-start structure. The parameter *r*_0_ = 20.0 Å controls the tolerance for distance deviations from the reference distance. The value of *Q* ranges from 0 to 1, with larger values indicating greater structural similarity to the compact two-start chromatin conformation.

We applied umbrella restraints using a harmonic potential with a spring constant of k = 150 *kJ/mol/*Å^2^. Umbrella windows were uniformly centered between *Q* = 0.30 and *Q* = 0.63, with an interval of 0.03. Each window was simulated for 20 million steps, and the first 10 million steps were excluded before constructing the unbiased free-energy profile. The simulation data were reweighted to reconstruct the unbiased free energy by using the Weighted Histogram Analysis Method (WHAM),^121^ as implemented in the SMOG software package.^66^

All 12-mer simulations were conducted at 300 K. To facilitate direct comparison with the sedimentation experiments performed at 20 ^◦^*C*,^68^ we reweighted the simulation ensembles to 20 ^◦^*C* before computing sedimentation coefficients (Figure S3).

#### Tuning DNA flexibility in Simulation

As in the DNA persistence-length simulation, DNA rigidity was modulated by varying the strength of the bonded interactions. Specifically, the bonded interaction scale, which rescales the force constants *k*_bond_, *k*_angle_, and *k*_fan bond_*_,n_* in Equations 10, 11, and 12, was varied around the baseline value of 0.8 to tune DNA flexibility. Additional details are provided in the Supplementary Material Section *DNA Persistence Length Parameterization*.

#### Charge Neutralization of Acetylated Histone Tail Sites

In all simulations, we modeled the acetylated lysine residues by neutralizing the positive charges of the corresponding residues in histone tails. Specifically, H3 tail acetylation was modeled by neutralizing the first seven lysine charges (K4, K9, K14, K27, K36, K37, and K56) on the H3 tail and the first four lysine residues (K5, K8, K12, and K20) on the H4 tail.

### Details of Simulation Analysis

#### Calculation of Sedimentation Coefficient for the 12-mer Chromatin Simulation

We used the HullRad method^122^ to calculate sedimentation coefficients for the 12-mer simulations. The method builds a convex hull based on the PDB structure of a biomolecule and calculates the sedimentation coefficient based on the estimated hydrodynamic volume using the following equation:

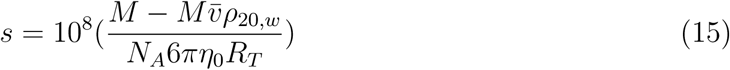

Here, *M* represents the molar mass of the molecule, *N_A_* is Avogadro’s number, and *v̄* is the partial specific volume. *ρ*_20_*_,w_* and *η*_0_ are the density and viscosity of water at 20^◦^*C*. *R_T_* is the translational hydrodynamic radius calculated from the convex hull of the target biomolecule. Further details of this calculation can be found in Fleming and Fleming ^122^. We adapted the code from Fleming and Fleming ^122^ for the coarse-grained 12-mer nucleosome array structures generated in our simulation.

#### Estimation of error bars

For sedimentation coefficients calculated from the 12-mer simulations, error bars were estimated as the standard deviation of the sedimentation coefficient, calculated using its probability distribution:

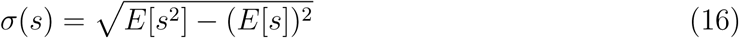

where *E*[*s*] is the expected value of the sedimentation coefficient *s*, calculated using Equation 15.

For all other simulations, error bars were estimated by dividing each trajectory into three equal-length segments and computing the corresponding quantity independently from each segment. The reported error bars were then calculated estimated as the standard deviation of these three independent estimates.

#### Determining Nucleosome Slab Densities

Nucleosome trajectories were analyzed using the MDTraj and MDAnalysis software pack-ages.^123–125^ To account for periodic boundary conditions, individual nucleosomes that were split across multiple periodic images were first reconstructed as whole molecules using the unwrap transformation implemented in make_mol_whole from openabc.utils.helper_functions. The center of mass (COM) for each nucleosome was then calculated based on the mass-weighted coordinates of its protein and DNA residues. To calculate condensate densities, we first identified the largest nucleosome cluster in each frame by using a graph-theoretic approach. Adjacent nucleosomes were identified using the FastNS grid search algorithm provided by MDAnalysis (Command: MDAnalysis.lib.nsgrid.FastNS), with a distance cutoff of 5.0 nm. These contacts were used to construct a proximity graph where nucleosomes were represented as nodes and contacts as edges. The largest condensate was defined as the largest connected component of this graph. To enable consistent spatial analysis across frames, the system was translated in each frame such that the center of the largest cluster was positioned at the center of the simulation box.

The nucleosome condensate density was computed by averaging the number density profile of simulated nucleosomes along the *z*-axis (the slab dimension). To ensure the analysis was performed on equilibrated simulations, the first half of each trajectory was excluded. The local number density, *ρ*(*z*), was calculated by binning the nucleosome COMs into slices of width Δ*z* = 10 nm according to the following equation:

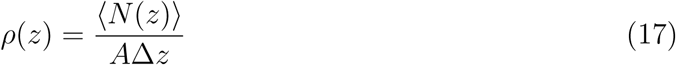

where ⟨*N* (*z*)⟩ is the average number of nucleosomes in the bin centered at position *z* and *A* is the cross-sectional area of the simulation box. All density values were converted to the unit of mM to facilitate comparison with experimental results.

#### Computation of Energetic Contributions from Nucleosomal Protein and DNA

To separate the contribution of protein-protein, protein-DNA, and DNA-DNA interactions during the simulations (Figure 3), we computed local interaction energies for individual nucleosomes within the condensate using the OpenMM simulation engine and the OpenABC framework. To isolate the interaction energy between a selected nucleosome and the rest of the system, we implemented an iterative relabeling procedure that distinguishes the selected nucleosome from all other nucleosomes. Specifically, for the *i*-th nucleosome, histone *α*-carbon beads were retyped from “CA” to “NP” and DNA beads were retyped from “DN” to “ND”. By defining custom interaction potentials that acts only on pairs involving these unique bead types (e.g., “NP”–“CA” for protein-protein interactions and ‘NP‘–‘DN‘ for protein–DNA interactions), we quantified protein-protein, protein-DNA, and DNA-DNA interactions between each individual nucleosome and the rest of the system throughout the simulation trajectory.

#### Computational of Inter-nucleosome Contacts

To quantify the physical connectivity of the nucleosome condensate, we analyzed internucleosome interactions mediated by disordered histone tails and nucleosomal DNA. These interactions were characterized using a distance-based contact analysis implemented with the MDAnalysis software package.^124,125^ For each nucleosome in the *N* = 100 system, a “contact” was defined when the distance between a bead in the specified histone-tail segments and a bead in the outer wrap of DNA from any other nucleosome fell below a screening-length-based cutoff, *d_c_*. Distances were calculated using np.linalg.norm. Under this unit-consistent framework, the physical cutoff distance was set equal to the Debye-Hűckel screening length:

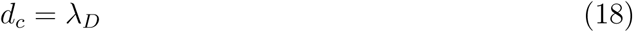

For each frame, we recorded the coordination number, *cn*, defined as the total number of histone tail–DNA bead pairs within the screening volume. To distinguish stable interactions from transient stochastic collisions, we applied a threshold of *θ* = 5, such that a stable inter-nucleosome interaction was assigned only when the number of contacts exceeded this value. Sensitivity analysis confirms that, although the quantitative count of inter-nucleosome contacts varied with *θ* ∈ {0, 5, 10, 15}, the qualitative salt-dependent trends remained robust (see Figure S9). Figure 3C describes the mean connectivity and fluctuations of the inter-nucleosome contacts across simulation trajectories, which were used to construct the connectivity networks shown in Figure 4.

#### Computation of Nucleosome Networks

To characterize the topological organization and connectivity of nucleosomes within the condensate, we performed graph-theoretic analysis using the networkx library. The system was represented as a bipartite graph *G* (Figure S10), in which each of the 100 nucleosomes was decomposed into two distinct node types: a histone protein node (*H_i_*) and a DNA node (*D_i_*), resulting in a total of 200 nodes.

Network edges were defined from the inter-nucleosomal contact maps. For each simulation frame, an edge was assigned between histone node *H_i_* and DNA node *D_j_* when larger-than-zero contacts were detected between the histone-tail residues of nucleosome *i* and the DNA nucleotides of nucleosome *j*. The resulting graph therefore describes the instantaneous physical connectivity among nucleosomes mediated by histone-tail–DNA interactions. To quantify the degree of physical aggregation, we identified the connected components of each graph using the connected_components function in the networkx library. The resulting cluster-size distribution was then used to calculate the weight-average component size, *S_w_*:

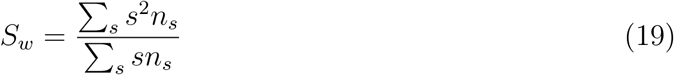

where *n_s_* is the number of clusters containing *s* nodes. This quantity represents the expected size of the cluster containing a randomly selected node.^126,127^

For each salt concentration, *S_w_* was calculated for each simulation frame and averaged over all frames to obtain the mean cluster size. The standard deviation across frames was also computed to quantify fluctuations in network connectivity. This analysis allowed us to track the system’s transition from a collection of small, isolated clusters to a large, interconnected percolating network, providing a quantitative measure of nucleosome aggregation as a function of salt concentration.

#### DNA Unwrapping

To monitor the structural integrity and conformational state of the nucleosomes within the condensate, we quantified the degree of DNA unwrapping as a function of salt concentration. DNA unwrapping was quantified as the average center-of-mass distance between the two terminal nucleotides of each nucleosomal DNA strand. Distance calculations were performed using the gmx distance tool from the GROMACS software package.^128^ For each of the *N* = 100 nucleosomes in the system, we defined index groups in custom index groups corresponding to the terminal beads of each DNA strand. Production trajectories across all salt concentrations were then processed by iterating over the nucleosome indices. An increase in the average distance between the two DNA ends indicates greater DNA unwrapping, allowing us to determine whether the high-density condensate environment or changes in salt concentration promote a more open nucleosomal conformation.

#### MSD, Diffusion Coefficient, and Viscosity Calculations

To characterize the rheological properties and dynamic behavior of the nucleosome condensate, we calculated the mean squared displacement (MSD), anomalous diffusion exponent, diffusion coefficient, and effective viscosity. The analysis was performed using a combination of the MDAnalysis and scipy libraries.

To ensure that the calculated displacement reflected internal condensate dynamics rather than global translational drift, we first applied a drift correction. A similar strategy was used in a recent experimental-computational study examining the effect of histone H3 tails on nucleosome condensate dynamics.^11^ For each frame *k*, the average velocity of the probe particles, **v**_avg_(*k*), was calculated as the mean displacement of all *N* nucleosomes between consecutive time steps:

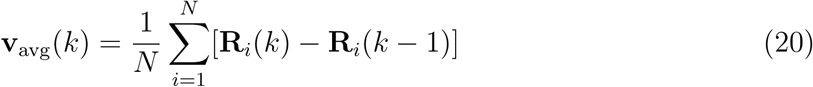

where *R_i_*(*k*) is the position of the *i*th nucleosome at frame *k*. The cumulative drift was then subtracted from the raw coordinates to obtain the drift-corrected coordinates:

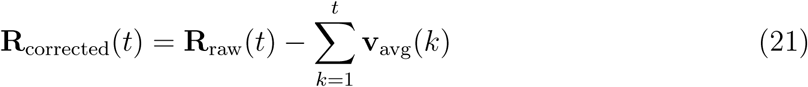

The ensemble-averaged MSD, ensemble_msd, was calculated as a function of lag time *τ*. For each probe particle, the MSD was defined as:

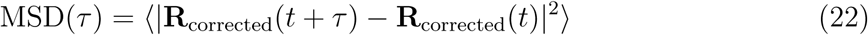

The resulting ensemble_msd was averaged over all nucleosomes in the system to improve statistical sampling.

To account for potentially anomalous diffusion, including sub-diffusive or super-diffusive behavior within the dense condensate, we fitted the ensemble_msd to a generalized power-law model using curve_fit:

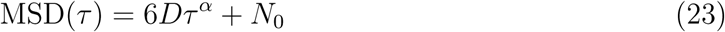

where *D* is the diffusion coefficient, *α* is the diffusion exponent (where *α* = 1 corresponds to Brownian motion), and *N*_0_ is a noise offset. These parameters were estimated by performing linear regression in log-log space using np.polyfit.

The effective viscosity (*η*) of the condensate was estimated from the fitted diffusion coefficient *D* using the Stokes-Einstein relation. After converting *D* from MDAnalysis units (Å^2^/ns) to SI units (m^2^*/*s), the viscosity was calculated as:

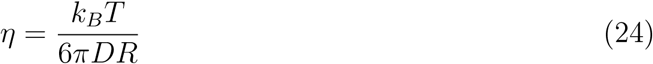

where *k_B_* is the Boltzmann constant, *T* is the simulation temperature of 300 K, and *R* is the effective hydrodynamic radius of the nucleosome (5.5 nm). The computed viscosity was reported in the unit of Pascal-seconds (Pa · s).

## Acknowledgement

This work was supported by startup funding from North Carolina State University and National Science Foundation (NSF) Grant MCB-2540505, with additional support from the NC State Genetics and Genomics Academy and the Comparative Medicine Institute. I.S. also acknowledges support from the NIH T32 Chemistry of Life Training Program at NC State University. We are grateful for valuable discussions with many experts, particularly Dr. Hong Wang, Dr. Michael Rubinstein, and Dr. Artem Rumyantsev.

## Data and Materials Availability

The implementation of the residue-resolution chromatin model, together with example workflows for setting up condensate simulations and performing post-simulation analyses, is available at our [GitHub repository](https://github.com/LinResearchGroup-NCSU/OpenABC_nucl).

## References

(1) Bertin, A.; Mangenot, S.; Renouard, M.; Durand, D.; Livolant, F. Structure and phase diagram of nucleosome core particles aggregated by multivalent cations. Biophys J 2007, 93, 3652–3663.

(2) Maeshima, K.; Rogge, R.; Tamura, S.; Joti, Y.; Hikima, T.; Szerlong, H.; Krause, C.; Herman, J.; Seidel, E.; DeLuca, J.; Ishikawa, T.; Hansen, J. C. Nucleosomal arrays self-assemble into supramolecular globular structures lacking 30-nm fibers. EMBO J 2016, 35, 1115–1132.

(3) Strom, A. R.; Emelyanov, A. V.; Mir, M.; Fyodorov, D. V.; Darzacq, X.; Karpen, G. H. Phase separation drives heterochromatin domain formation. Nature 2017, 547, 241–245.

(4) Gibson, B. A.; Doolittle, L. K.; Schneider, M. W.; Jensen, L. E.; Gamarra, N.; Henry, L.; Gerlich, D. W.; Redding, S.; Rosen, M. K. Organization of Chromatin by Intrinsic and Regulated Phase Separation. Cell 2019, 179, 470–484.e21.

(5) Hammonds, E. F.; Harwig, M. C.; Paintsil, E. A.; Tillison, E. A.; Hill, R. B.; Morrison, E. A. Histone H3 and H4 tails play an important role in nucleosome phase separation. Biophysical Chemistry 2022, 283, 106767.

(6) Farr, S. E.; Woods, E. J.; Joseph, J. A.; Garaizar, A.; Collepardo-Guevara, R. Nucleosome plasticity is a critical element of chromatin liquid–liquid phase separation and multivalent nucleosome interactions. Nat Commun 2021, 12, 2883.

(7) Zhou, H.; Hutchings, J.; Shiozaki, M.; Zhao, X.; Doolittle, L. K.; Yang, S.; Yan, R.; Jean, N.; Riggi, M.; Yu, Z.; Villa, E.; Rosen, M. K. Quantitative spatial analysis of chromatin biomolecular condensates using cryoelectron tomography. Proc Natl Acad Sci U S A 2025, 122, e2426449122.

(8) Zhou, H. et al. Multiscale structure of chromatin condensates explains phase separation and material properties. Science 2025, 390, eadv6588.

(9) Fujishiro, S.; Sasai, M.; Maeshima, K. Chromatin domains in the cell: Phase separation and condensation. Current Opinion in Structural Biology 2025, 91, 103006.

(10) Qiu, Y.; Liu, S.; Lin, X.; Unarta, I. C.; Huang, X.; Zhang, B. Nucleosome condensate and linker DNA alter chromatin folding pathways and rates. Biophysical Journal 2026, 125, 282–293.

(11) Hammonds, E. F.; Singh, A.; Suresh, K. K.; Yang, S.; Meidl Zahorodny, S. S.; Gupta, R.; Potoyan, D. A.; Banerjee, P. R.; Morrison, E. A. Histone H3 tail charge patterns govern nucleosome condensate formation and dynamics. Nucleic Acids Research 2026, 54, gkag050.

(12) Lieberman-Aiden, E. et al. Comprehensive Mapping of Long-Range Interactions Reveals Folding Principles of the Human Genome. Science 2009, 326, 289–293.

(13) Rao, S. S.; Huntley, M. H.; Durand, N. C.; Stamenova, E. K.; Bochkov, I. D.; Robinson, J. T.; Sanborn, A. L.; Machol, I.; Omer, A. D.; Lander, E. S.; Aiden, E. L. A 3D Map of the Human Genome at Kilobase Resolution Reveals Principles of Chromatin Looping. Cell 2014, 159, 1665–1680.

(14) Goel, V. Y.; Huseyin, M. K.; Hansen, A. S. Region Capture Micro-C reveals coalescence of enhancers and promoters into nested microcompartments. Nat Genet 2023, 55, 1048–1056.

(15) Oberbeckmann, E.; Quililan, K.; Cramer, P.; Oudelaar, A. M. In vitro reconstitution of chromatin domains shows a role for nucleosome positioning in 3D genome organization. Nat Genet 2024, 56, 483–492.

(16) Hsieh, T.-H. S.; Cattoglio, C.; Slobodyanyuk, E.; Hansen, A. S.; Rando, O. J.; Tjian, R.; Darzacq, X. Resolving the 3D Landscape of Transcription-Linked Mammalian Chromatin Folding. Mol Cell 2020, 78, 539–553.e8.

(17) Nozaki, T.; Shinkai, S.; Ide, S.; Higashi, K.; Tamura, S.; Shimazoe, M. A.; Nakagawa, M.; Suzuki, Y.; Okada, Y.; Sasai, M.; Onami, S.; Kurokawa, K.; Iida, S.; Maeshima, K. Condensed but liquid-like domain organization of active chromatin regions in living human cells. Sci Adv 2023, 9, eadf1488.

(18) Bintu, B.; Mateo, L. J.; Su, J.-H.; Sinnott-Armstrong, N. A.; Parker, M.; Kinrot, S.; Yamaya, K.; Boettiger, A. N.; Zhuang, X. Super-resolution chromatin tracing reveals domains and cooperative interactions in single cells. Science 2018, 362, eaau1783.

(19) Miron, E.; Oldenkamp, R.; Brown, J. M.; Pinto, D. M. S.; Xu, C. S.; Faria, A. R.; Shaban, H. A.; Rhodes, J. D. P.; Innocent, C.; de Ornellas, S.; Hess, H. F.; Buckle, V.; Schermelleh, L. Chromatin arranges in chains of mesoscale domains with nanoscale functional topography independent of cohesin. Sci Adv 2020, 6, eaba8811.

(20) Boettiger, A. N.; Bintu, B.; Moffitt, J. R.; Wang, S.; Beliveau, B. J.; Fudenberg, G.; Imakaev, M.; Mirny, L. A.; Wu, C.-t.; Zhuang, X. Super-resolution imaging reveals distinct chromatin folding for different epigenetic states. Nature 2016, 529, 418–422.

(21) Iida, S.; Ide, S.; Tamura, S.; Sasai, M.; Tani, T.; Goto, T.; Shribak, M.; Maeshima, K. Orientation-independent-DIC imaging reveals that a transient rise in depletion attraction contributes to mitotic chromosome condensation. Proc Natl Acad Sci U S A 2024, 121, e2403153121.

(22) Jost, D.; Carrivain, P.; Cavalli, G.; Vaillant, C. Modeling epigenome folding: formation and dynamics of topologically associated chromatin domains. Nucleic Acids Res 2014, 42, 9553–9561.

(23) Jiang, Z.; Qi, Y.; Kamat, K.; Zhang, B. Phase Separation and Correlated Motions in Motorized Genome. J Phys Chem B 2022, 126, 5619–5628.

(24) Shin, S.; Shi, G.; Cho, H. W.; Thirumalai, D. Transcription-induced active forces suppress chromatin motion. Proc Natl Acad Sci U S A 2024, 121, e2307309121.

(25) Brahmachari, S.; Markovich, T.; MacKintosh, F. C.; Onuchic, J. N. Temporally Correlated Active Forces Drive Segregation and Enhanced Dynamics in Chromosome Polymers. PRX Life 2024, 2, 033003.

(26) Strickfaden, H.; Tolsma, T. O.; Sharma, A.; Underhill, D. A.; Hansen, J. C.; Hendzel, M. J. Condensed Chromatin Behaves like a Solid on the Mesoscale In Vitro and in Living Cells. Cell 2020, 183, 1772–1784.e13.

(27) Eshghi, I.; Eaton, J. A.; Zidovska, A. Interphase Chromatin Undergoes a Local Sol-Gel Transition upon Cell Differentiation. Phys. Rev. Lett. 2021, 126, 228101.

(28) Chen, Q.; Zhao, L.; Soman, A.; Arkhipova, A. Y.; Li, J.; Li, H.; Chen, Y.; Shi, X.; Nordenskiöld, L. Chromatin Liquid–Liquid Phase Separation (LLPS) Is Regulated by Ionic Conditions and Fiber Length. Cells 2022, 11, 3145.

(29) Park, S.; Merino-Urteaga, R.; Karwacki-Neisius, V.; Carrizo, G. E.; Athreya, A.; Marin-Gonzalez, A.; Benning, N. A.; Park, J.; Mitchener, M. M.; Bhanu, N. V.; Garcia, B. A.; Zhang, B.; Muir, T. W.; Pearce, E. L.; Ha, T. Native nucleosomes intrinsically encode genome organization principles. Nature 2025, 643, 572–581.

(30) Kent, S.; Brown, K.; Yang, C.-h.; Alsaihati, N.; Tian, C.; Wang, H.; Ren, X. Phase-Separated Transcriptional Condensates Accelerate Target-Search Process Revealed by Live-Cell Single-Molecule Imaging. Cell Reports 2020, 33, 108248.

(31) Eeftens, J. M.; Kapoor, M.; Michieletto, D.; Brangwynne, C. P. Polycomb conden-sates can promote epigenetic marks but are not required for sustained chromatin compaction. Nat Commun 2021, 12, 5888.

(32) Seif, E.; Kang, J. J.; Sasseville, C.; Senkovich, O.; Kaltashov, A.; Boulier, E. L.; Kapur, I.; Kim, C. A.; Francis, N. J. Phase separation by the polyhomeotic sterile alpha motif compartmentalizes Polycomb Group proteins and enhances their activity. Nat Commun 2020, 11, 5609.

(33) Larson, A. G.; Elnatan, D.; Keenen, M. M.; Trnka, M. J.; Johnston, J. B.; Burlingame, A. L.; Agard, D. A.; Redding, S.; Narlikar, G. J. Liquid droplet formation by HP1 suggests a role for phase separation in heterochromatin. Nature 2017, 547, 236–240.

(34) Maeshima, K.; Matsuda, T.; Shindo, Y.; Imamura, H.; Tamura, S.; Imai, R.; Kawakami, S.; Nagashima, R.; Soga, T.; Noji, H.; Oka, K.; Nagai, T. A Transient Rise in Free Mg2+ Ions Released from ATP-Mg Hydrolysis Contributes to Mitotic Chromosome Condensation. Curr Biol 2018, 28, 444–451.e6.

(35) Nott, T.; Petsalaki, E.; Farber, P.; Jervis, D.; Fussner, E.; Plochowietz, A.; Craggs, T. D.; Bazett-Jones, D.; Pawson, T.; Forman-Kay, J.; Baldwin, A. Phase Transition of a Disordered Nuage Protein Generates Environmentally Responsive Membraneless Organelles. Molecular Cell 2015, 57, 936–947.

(36) Lin, Y.-H.; Forman-Kay, J. D.; Chan, H. S. Sequence-Specific Polyampholyte Phase Separation in Membraneless Organelles. Phys Rev Lett 2016, 117, 178101.

(37) Arya, G.; Schlick, T. Role of histone tails in chromatin folding revealed by a mesoscopic oligonucleosome model. Proc. Natl. Acad. Sci. U.S.A. 2006, 103, 16236–16241.

(38) Fan, Y.; Korolev, N.; Lyubartsev, A. P.; Nordenskiöld, L. An advanced coarse-grained nucleosome core particle model for computer simulations of nucleosome-nucleosome interactions under varying ionic conditions. PLoS One 2013, 8, e54228.

(39) Lin, X.; Zhang, B. Explicit ion modeling predicts physicochemical interactions for chromatin organization. eLife 2024, 12, RP90073.

(40) Rumyantsev, A. M.; Gavrilov, A. A.; Johner, A. Complete Diagram of Conformational Regimes for Polyampholytic Disordered Proteins. Macromolecules 2024, 57, 5533–5544.

(41) Bowman, W. A.; Rubinstein, M.; Tan, J. S. PolyelectrolyteGelatin Complexation: Light-Scattering Study. Macromolecules 1997, 30, 3262–3270.

(42) Strahl, B. D.; Allis, C. D. The language of covalent histone modifications. Nature 2000, 403, 41–45.

(43) Baylin, S. B.; Jones, P. A. A decade of exploring the cancer epigenome - biological and translational implications. Nat Rev Cancer 2011, 11, 726–734.

(44) Allis, C. D.; Jenuwein, T. The molecular hallmarks of epigenetic control. Nat Rev Genet 2016, 17, 487–500.

(45) Brower-Toland, B.; Wacker, D. A.; Fulbright, R. M.; Lis, J. T.; Kraus, W. L.; Wang, M. D. Specific Contributions of Histone Tails and their Acetylation to the Mechanical Stability of Nucleosomes. Journal of Molecular Biology 2005, 346, 135–146.

(46) Zhang, R.; Erler, J.; Langowski, J. Histone Acetylation Regulates Chromatin Accessibility: Role of H4K16 in Inter-nucleosome Interaction. Biophys J 2017, 112, 450–459.

(47) Kim, T.; Nosella, M.; Bolik-Coulon, N.; Harkness, R.; Huang, S.; Kay, L. Correlating histone acetylation with nucleosome core particle dynamics and function. Proc. Natl. Acad. Sci. U.S.A. 2023, 120, e2301063120.

(48) Shogren-Knaak, M.; Ishii, H.; Sun, J.-M.; Pazin, M. J.; Davie, J. R.; Peterson, C. L. Histone H4-K16 acetylation controls chromatin structure and protein interactions. Science 2006, 311, 844–847.

(49) Li, R.; Lin, X. Connected Chromatin Amplifies Acetylation-Modulated Nucleosome Interactions. Biochemistry 2025, 64, 1222–1232.

(50) Shia, W.-J.; Pattenden, S. G.; Workman, J. L. Histone H4 lysine 16 acetylation breaks the genome’s silence. Genome Biol 2006, 7, 217.

(51) Nitsch, S. et al. H4K16 acylations destabilize chromatin architecture and facilitate transcriptional response during metabolic perturbations. Mol Cell 2026, 86, 24–40.e10.

(52) Wedemann, G.; Langowski, J. Computer simulation of the 30-nanometer chromatin fiber. Biophys J 2002, 82, 2847–2859.

(53) Kepper, N.; Foethke, D.; Stehr, R.; Wedemann, G.; Rippe, K. Nucleosome geometry and internucleosomal interactions control the chromatin fiber conformation. Biophys J 2008, 95, 3692–3705.

(54) D. Bascom, G.; Schlick, T. Nuclear Architecture and Dynamics; Elsevier, 2018; pp 123–147.

(55) Golembeski, A.; Lequieu, J. A Molecular View into the Structure and Dynamics of Phase-Separated Chromatin. J Phys Chem B 2024, 128, 10593–10603.

(56) Peng, Y.; Li, S.; Onufriev, A.; Landsman, D.; Panchenko, A. R. Binding of regulatory proteins to nucleosomes is modulated by dynamic histone tails. Nat Commun 2021, 12, 5280.

(57) Armeev, G. A.; Kniazeva, A. S.; Komarova, G. A.; Kirpichnikov, M. P.; Shaytan, A. K. Histone dynamics mediate DNA unwrapping and sliding in nucleosomes. Nat Commun 2021, 12, 2387.

(58) Izadi, S.; Anandakrishnan, R.; Onufriev, A. V. Implicit Solvent Model for Million-Atom Atomistic Simulations: Insights into the Organization of 30-nm Chromatin Fiber. J Chem Theory Comput 2016, 12, 5946–5959.

(59) Woods, D. C.; Rodŕıguez-Ropero, F.; Wereszczynski, J. The Dynamic Influence of Linker Histone Saturation within the Poly-Nucleosome Array. J Mol Biol 2021, 433, 166902.

(60) Russell, K.; Chen, Y.; Espinosa, J. R.; Gil, D. F.; Zhou, H.; Maristany, M. J.; Perez-Lopez, J. I.; Huertas, J.; Orozco, M.; Rosen, M. K.; Collepardo-Guevara, R. Near-atomistic simulations reveal the molecular principles that control chromatin structure and phase separation. bioRxiv 2025, 2025.11.17.688899.

(61) Huertas, J.; Woods, E. J.; Collepardo-Guevara, R. Multiscale modelling of chromatin organisation: Resolving nucleosomes at near-atomistic resolution inside genes. Curr Opin Cell Biol 2022, 75, 102067.

(62) Maristany, M. J.; Emelianova, A.; Chew, P. Y.; Aguirre, A.; Collepardo-Guevara, R.; Joseph, J. A. Modeling biomolecular condensates across scales: Atomistic, coarse-grained, and data-driven approaches. Adv Phys X 2025, 10, 2592547.

(63) Liu, S.; Wang, C.; Latham, A. P.; Ding, X.; Zhang, B. OpenABC enables flexible, simplified, and efficient GPU accelerated simulations of biomolecular condensates. PLoS Comput Biol 2023, 19, e1011442.

(64) Eastman, P. et al. OpenMM 8: Molecular Dynamics Simulation with Machine Learning Potentials. J. Phys. Chem. B 2024, 128, 109–116.

(65) Clementi, C.; Nymeyer, H.; Onuchic, J. N. Topological and energetic factors: what determines the structural details of the transition state ensemble and “en-route” intermediates for protein folding? an investigation for small globular proteins. Journal of Molecular Biology 2000, 298, 937–953.

(66) Noel, J. K.; Levi, M.; Raghunathan, M.; Lammert, H.; Hayes, R. L.; Onuchic, J. N.; Whitford, P. C. SMOG 2: A Versatile Software Package for Generating Structure-Based Models. PLoS Comput Biol 2016, 12, e1004794.

(67) Savelyev, A.; Papoian, G. A. Chemically accurate coarse graining of double-stranded DNA. Proc. Natl. Acad. Sci. U.S.A. 2010, 107, 20340–20345.

(68) Correll, S. J.; Schubert, M. H.; Grigoryev, S. A. Short nucleosome repeats impose rotational modulations on chromatin fibre folding: Rotational modulations of chromatin folding. The EMBO Journal 2012, 31, 2416–2426.

(69) Gao, J.; Li, H.; Tan, S.; Zhou, R.; Lee, T.-H. Roles of histone chaperone Nap1 and histone acetylation in regulating phase-separation of nucleosome arrays. Nat Commun 2025, 16, 10672.

(70) Korolev, N.; Allahverdi, A.; Yang, Y.; Fan, Y.; Lyubartsev, A. P.; Nordenskiöld, L. Electrostatic origin of salt-induced nucleosome array compaction. Biophys J 2010, 99, 1896–1905.

(71) Gan, H. H.; Schlick, T. Chromatin ionic atmosphere analyzed by a mesoscale electrostatic approach. Biophys J 2010, 99, 2587–2596.

(72) Dignon, G. L.; Zheng, W.; Kim, Y. C.; Best, R. B.; Mittal, J. Sequence determinants of protein phase behavior from a coarse-grained model. PLoS Comput Biol 2018, 14, e1005941.

(73) Latham, A. P.; Zhang, B. On the stability and layered organization of protein-DNA condensates. Biophys J 2022, 121, 1727–1737.

(74) Ricci, M. A.; Manzo, C.; Garćıa-Parajo, M. F.; Lakadamyali, M.; Cosma, M. P. Chromatin fibers are formed by heterogeneous groups of nucleosomes in vivo. Cell 2015, 160, 1145–1158.

(75) Portillo-Ledesma, S.; Tsao, L. H.; Wagley, M.; Lakadamyali, M.; Cosma, M. P.; Schlick, T. Nucleosome Clutches are Regulated by Chromatin Internal Parameters. J Mol Biol 2021, 433, 166701.

(76) Liu, S.; Lin, X.; Zhang, B. Chromatin fiber breaks into clutches under tension and crowding. Nucleic Acids Res 2022, 50, 9738–9747.

(77) Farag, M.; Cohen, S. R.; Borcherds, W. M.; Bremer, A.; Mittag, T.; Pappu, R. V. Condensates formed by prion-like low-complexity domains have small-world network structures and interfaces defined by expanded conformations. Nat Commun 2022, 13, 7722.

(78) Grigorev, V.; Wingreen, N. S.; Zhang, Y. Conformational Entropy of Intrinsically Disordered Proteins Bars Intruders from Biomolecular Condensates. PRX Life 2025, 3, 013011.

(79) Geggier, S.; Vologodskii, A. Sequence dependence of DNA bending rigidity. Proc Natl Acad Sci U S A 2010, 107, 15421–15426.

(80) Ortiz, V.; de Pablo, J. J. Molecular origins of DNA flexibility: sequence effects on conformational and mechanical properties. Phys Rev Lett 2011, 106, 238107.

(81) Lin, X.; Leicher, R.; Liu, S.; Zhang, B. Cooperative DNA looping by PRC2 complexes. Nucleic Acids Research 2021, 49, 6238–6248.

(82) Farŕe-Gil, D.; Arcon, J. P.; Laughton, C. A.; Orozco, M. CGeNArate: a sequence-dependent coarse-grained model of DNA for accurate atomistic MD simulations of kb-long duplexes. Nucleic Acids Research 2024, 52, 6791–6801.

(83) Fenley, A. T.; Anandakrishnan, R.; Kidane, Y. H.; Onufriev, A. V. Modulation of nucleosomal DNA accessibility via charge-altering post-translational modifications in histone core. Epigenetics & Chromatin 2018, 11, 11.

(84) Zheng, C.; Lu, X.; Hansen, J. C.; Hayes, J. J. Salt-dependent intra- and internucleosomal interactions of the H3 tail domain in a model oligonucleosomal array. J Biol Chem 2005, 280, 33552–33557.

(85) Bascom, G. D.; Schlick, T. Chromatin Fiber Folding Directed by Cooperative Histone Tail Acetylation and Linker Histone Binding. Biophys J 2018, 114, 2376–2385.

(86) Takada, S.; Kanada, R.; Tan, C.; Terakawa, T.; Li, W.; Kenzaki, H. Modeling Structural Dynamics of Biomolecular Complexes by Coarse-Grained Molecular Simulations. Acc. Chem. Res. 2015, 48, 3026–3035.

(87) Stradner, A.; Sedgwick, H.; Cardinaux, F.; Poon, W. C. K.; Egelhaaf, S. U.; Schurtenberger, P. Equilibrium cluster formation in concentrated protein solutions and colloids. Nature 2004, 432, 492–495.

(88) Zhang, M.; Díaz-Celis, C.; Onoa, B.; Cañari-Chumpitaz, C.; Requejo, K. I.; Liu, J.; Vien, M.; Nogales, E.; Ren, G.; Bustamante, C. Molecular organization of the early stages of nucleosome phase separation visualized by cryo-electron tomography. Molecular Cell 2022, 82, 3000–3014.e9.

(89) Kar, M.; Dar, F.; Welsh, T. J.; Vogel, L. T.; Kűhnemuth, R.; Majumdar, A.; Krainer, G.; Franzmann, T. M.; Alberti, S.; Seidel, C. A. M.; Knowles, T. P. J.; Hyman, A. A.; Pappu, R. V. Phase-separating RNA-binding proteins form heterogeneous distributions of clusters in subsaturated solutions. Proc Natl Acad Sci U S A 2022, 119, e2202222119.

(90) Mittag, T.; Pappu, R. V. A conceptual framework for understanding phase separation and addressing open questions and challenges. Mol Cell 2022, 82, 2201–2214.

(91) Maclennan, J.; Seul, M. Novel stripe textures in nonchiral hexatic liquid-crystal films. Phys. Rev. Lett. 1992, 69, 2082–2085.

(92) Seul, M.; Andelman, D. Domain Shapes and Patterns: The Phenomenology of Modulated Phases. Science 1995, 267, 476–483.

(93) Banani, S. F.; Lee, H. O.; Hyman, A. A.; Rosen, M. K. Biomolecular condensates: organizers of cellular biochemistry. Nat Rev Mol Cell Biol 2017, 18, 285–298.

(94) Zhang, M.; Díaz-Celis, C.; Liu, J.; Tao, J.; Ashby, P. D.; Bustamante, C.; Ren, G. Angle between DNA linker and nucleosome core particle regulates array compaction revealed by individual-particle cryo-electron tomography. Nat Commun 2024, 15, 4395.

(95) Chakraborty, D.; Mondal, B.; Thirumalai, D. Brewing COFFEE: A Sequence-Specific Coarse-Grained Energy Function for Simulations of DNAProtein Complexes. J. Chem. Theory Comput. 2024, 20, 1398–1413.

(96) Zhang, Y.; Silvernail, I.; Lin, Z.; Lin, X. Interpretable protein-DNA interactions captured by structure-sequence optimization. Elife 2025, 14, RP105565.

(97) Tesei, G.; Lindorff-Larsen, K. Improved predictions of phase behaviour of intrinsically disordered proteins by tuning the interaction range. Open Res Europe 2022, 2, 94.

(98) R. Tejedor, A.; Aguirre Gonzalez, A.; Maristany, M. J.; Chew, P. Y.; Russell, K.; Ramirez, J.; Espinosa, J. R.; Collepardo-Guevara, R. Chemically Informed Coarse-Graining of Electrostatic Forces in Charge-Rich Biomolecular Condensates. ACS Cent. Sci. 2025, 11, 302–321.

(99) Riveros, I.; Zhang, B. NEAT-DNA: A Chemically Accurate, Sequence-Dependent Coarse-Grained Model for Large-Scale DNA Simulations. bioRxiv 2025, 2025.11.07.687145.

(100) Farag, M.; Borcherds, W. M.; Bremer, A.; Mittag, T.; Pappu, R. V. Phase separation of protein mixtures is driven by the interplay of homotypic and heterotypic interactions. Nat Commun 2023, 14, 5527.

(101) Wang, J.; Choi, J.-M.; Holehouse, A. S.; Lee, H. O.; Zhang, X.; Jahnel, M.; Maharana, S.; Lemaitre, R.; Pozniakovsky, A.; Drechsel, D.; Poser, I.; Pappu, R. V.; Alberti, S.; Hyman, A. A. A Molecular Grammar Governing the Driving Forces for Phase Separation of Prion-like RNA Binding Proteins. Cell 2018, 174, 688–699.e16.

(102) Langstein-Skora, I. et al. Sequence and chemical specificity define the functional landscape of intrinsically disordered regions. Nat Cell Biol 2026, 28, 323–337.

(103) Latham, A. P.; Zhang, B. Consistent Force Field Captures Homologue-Resolved HP1 Phase Separation. J Chem Theory Comput 2021, 17, 3134–3144.

(104) Tesei, G.; Schulze, T. K.; Crehuet, R.; Lindorff-Larsen, K. Accurate model of liquid-liquid phase behavior of intrinsically disordered proteins from optimization of single-chain properties. Proc Natl Acad Sci U S A 2021, 118, e2111696118.

(105) Joseph, J. A.; Reinhardt, A.; Aguirre, A.; Chew, P. Y.; Russell, K. O.; Espinosa, J. R.; Garaizar, A.; Collepardo-Guevara, R. Physics-driven coarse-grained model for biomolecular phase separation with near-quantitative accuracy. Nat Comput Sci 2021, 1, 732–743.

(106) Thornton, T.; Lin, X. Efficient RNA Folding Simulation via a Structure-Based Single-Site-Per-Nucleotide Model. 2025; http://biorxiv.org/lookup/doi/ 10.64898/2025.12.13.694107.

(107) Leicher, R.; Osunsade, A.; Chua, G. N. L.; Faulkner, S. C.; Latham, A. P.; Wat-ters, J. W.; Nguyen, T.; Beckwitt, E. C.; Christodoulou-Rubalcava, S.; Young, P. G.; Zhang, B.; David, Y.; Liu, S. Single-stranded nucleic acid binding and coacervation by linker histone H1. Nat Struct Mol Biol 2022, 29, 463–471.

(108) Savelyev, A.; Papoian, G. A. Molecular renormalization group coarse-graining of polymer chains: application to double-stranded DNA. Biophys J 2009, 96, 4044–4052.

(109) Latham, A. P.; Zhang, B. Maximum Entropy Optimized Force Field for Intrinsically Disordered Proteins. J Chem Theory Comput 2020, 16, 773–781.

(110) Noel, J. K.; Whitford, P. C.; Onuchic, J. N. The shadow map: a general contact definition for capturing the dynamics of biomolecular folding and function. J Phys Chem B 2012, 116, 8692–8702.

(111) Ding, X.; Lin, X.; Zhang, B. Stability and folding pathways of tetra-nucleosome from six-dimensional free energy surface. Nat Commun 2021, 12, 1091.

(112) Davey, C. A.; Sargent, D. F.; Luger, K.; Maeder, A. W.; Richmond, T. J. Solvent mediated interactions in the structure of the nucleosome core particle at 1.9 a resolution. J Mol Biol 2002, 319, 1097–1113.

(113) Vasudevan, D.; Chua, E. Y.; Davey, C. A. Crystal Structures of Nucleosome Core Particles Containing the ‘601’ Strong Positioning Sequence. Journal of Molecular Biology 2010, 403, 1–10.

(114) Schalch, T.; Duda, S.; Sargent, D. F.; Richmond, T. J. X-ray structure of a tetranucleosome and its implications for the chromatin fibre. Nature 2005, 436, 138–141.

(115) Lu, X.-J.; Olson, W. K. 3DNA: a software package for the analysis, rebuilding and visualization of three-dimensional nucleic acid structures. Nucleic Acids Res 2003, 31, 5108–5121.

(116) Zhang, Z.; Liu, X.; Yan, K.; Tuckerman, M. E.; Liu, J. Unified Efficient Thermostat Scheme for the Canonical Ensemble with Holonomic or Isokinetic Constraints via Molecular Dynamics. J Phys Chem A 2019, 123, 6056–6079.

(117) Torrie, G.; Valleau, J. Nonphysical sampling distributions in Monte Carlo free-energy estimation: Umbrella sampling. Journal of Computational Physics 1977, 23, 187–199.

(118) The PLUMED consortium Promoting transparency and reproducibility in enhanced molecular simulations. Nat Methods 2019, 16, 670–673.

(119) Moller, J.; Lequieu, J.; De Pablo, J. J. The Free Energy Landscape of Internucleosome Interactions and Its Relation to Chromatin Fiber Structure. ACS Cent. Sci. 2019, 5, 341–348.

(120) Song, F.; Chen, P.; Sun, D.; Wang, M.; Dong, L.; Liang, D.; Xu, R.-M.; Zhu, P.; Li, G. Cryo-EM study of the chromatin fiber reveals a double helix twisted by tetranucleosomal units. Science 2014, 344, 376–380.

(121) Kumar, S.; Rosenberg, J. M.; Bouzida, D.; Swendsen, R. H.; Kollman, P. A. THE weighted histogram analysis method for free-energy calculations on biomolecules. I. The method. J Comput Chem 1992, 13, 1011–1021.

(122) Fleming, P. J.; Fleming, K. G. HullRad: Fast Calculations of Folded and Disordered Protein and Nucleic Acid Hydrodynamic Properties. Biophys J 2018, 114, 856–869.

(123) McGibbon, R. T.; Beauchamp, K. A.; Harrigan, M. P.; Klein, C.; Swails, J. M.; Hernández, C. X.; Schwantes, C. R.; Wang, L.-P.; Lane, T. J.; Pande, V. S. MDTraj: A Modern Open Library for the Analysis of Molecular Dynamics Trajectories. Biophys J 2015, 109, 1528–1532.

(124) Gowers, R.; Linke, M.; Barnoud, J.; Reddy, T.; Melo, M.; Seyler, S.; Domański, J.; Dotson, D.; Buchoux, S.; Kenney, I.; Beckstein, O. MDAnalysis: A Python Package for the Rapid Analysis of Molecular Dynamics Simulations. Austin, Texas, 2016; pp 98–105.

(125) Michaud-Agrawal, N.; Denning, E. J.; Woolf, T. B.; Beckstein, O. MDAnalysis: a toolkit for the analysis of molecular dynamics simulations. J Comput Chem 2011, 32, 2319–2327.

(126) Hassan, M. K.; Rahman, M. M. Percolation on a multifractal scale-free planar stochastic lattice and its universality class. *Phys*. Rev. E 2015, 92, 040101.

(127) Li, M.; Liu, R.-R.; Lű, L.; Hu, M.-B.; Xu, S.; Zhang, Y.-C. Percolation on complex networks: Theory and application. Physics Reports 2021, 907, 1–68.

(128) Abraham, M. J.; Murtola, T.; Schulz, R.; Páll, S.; Smith, J. C.; Hess, B.; Lindahl, E. GROMACS: High performance molecular simulations through multi-level parallelism from laptops to supercomputers. SoftwareX 2015, 1-2, 19–25.

