## Supplementary Information for "Charge Imbalance Drives Salt-Optimized Nucleosome Phase Separation under Physiological Conditions"

### Contents

**1 Figures** **S-3**

1.1 Model Validation . . . . . . . . . . S-3

1.1.1 DNA Persistence Length . . . . . . . . . . S-3

1.1.2 Inter-nucleosome Interactions . . . . . . . . . . S-4

1.1.3 12-mer Nucleosomal Array Structure . . . . . . . . . . S-5

1.2 Benchmark of Model Efficiency . . . . . . . . . . . . . . . . . . . . . . . . . S-5

1.3 Determining Dense and Dilute Phases . . . . . . . . . . . . . . . . . . . . . . . S-8

1.4 Calculating Critical Concentrations . . . . . . . . . . . . . . . . . . . . . . . . S-9

1.5 Cleaning the Data for phase diagram . . . . . . . . . . . . . . . . . . . . . . . S-9

1.6 Additional Simulation Analyses . . . . . . . . . . . . . . . . . . . . . . . . . S-13

1.6.1 Definition of Nucleosome-Nucleosome Contact Thresholds . . . . . . S-13

1.6.2 Spatiotemporal Analysis of Nucleosome Interaction Networks . . . . . S-14

1.6.3 Mechanical and Chemical Modulation of Condensate Architecture . . S-16

### 1 Figures

#### 1.1 Model Validation

##### 1.1.1 DNA Persistence Length

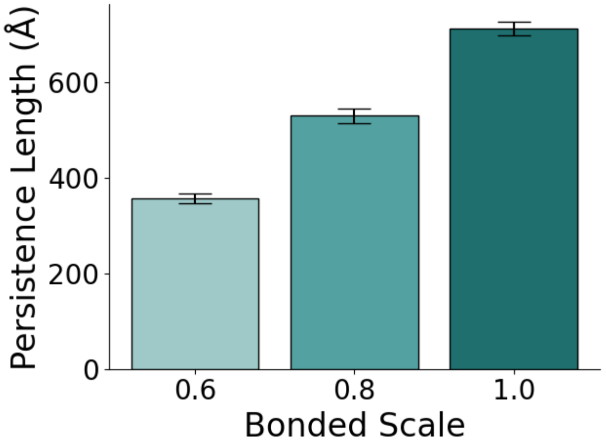


Figure S1: **Parameterization and validation of DNA persistence length.** Calculated DNA persistence length ($\AA$) as a function of the model's bending rigidity parameter, with bonded scale values of 0.6, 0.8, and 1.0. The close agreement between the model prediction at the bonded scale of 0.8 and the experimentally measured persistence length of approximately 500 $\AA$ for a 150-bp DNA segment validates the mechanical accuracy of the baseline model, establishing a rigorous benchmark for comparing chromatin with mechanically flexible (0.6), baseline (0.8), and more rigid (1.0) DNA.

To ensure that the coarse-grained DNA model accurately captures the mechanical properties of B-DNA, we compared the simulated DNA persistence length ($L_p$) with the experimentally reported values.[S1–S3] DNA was modeled using the molecular renormalization group (MRG) model, where the internal stiffness of DNA is controlled by the parameter `bonded_energy_scale`. The persistence length was determined by simulating a standalone 150-base-pair (bp) DNA segment. We varied the global scaling factor applied to the bonded DNA interactions, including $k_{\text{bond}}$, $k_{\text{angle}}$, and $k_{\text{fan-bond}}$ (referred to as "bonded scale") and calculated the resulting $L_p$ values. A bonded scale of 0.8 reproduced the canonical experimental persistence length of double-stranded DNA, $L_p \approx 50\,\text{nm}$[S1–S3] (Figure S1). We therefore used a bonded scale of 0.8 as the baseline DNA stiffness in subsequent simulations.

All calibration simulations were performed using the OpenMM molecular dynamics engine[S4] and used the same parameters as the production simulations (see *Methods and Materials* Section), except that the friction coefficient of 1 $\mathrm{ps}^{-1}$ was used for the Langevin Middle Integrator. The friction coefficient does not affect the determination of the equilibrium DNA persistence length from the simulation.

##### 1.1.2 Inter-nucleosome Interactions

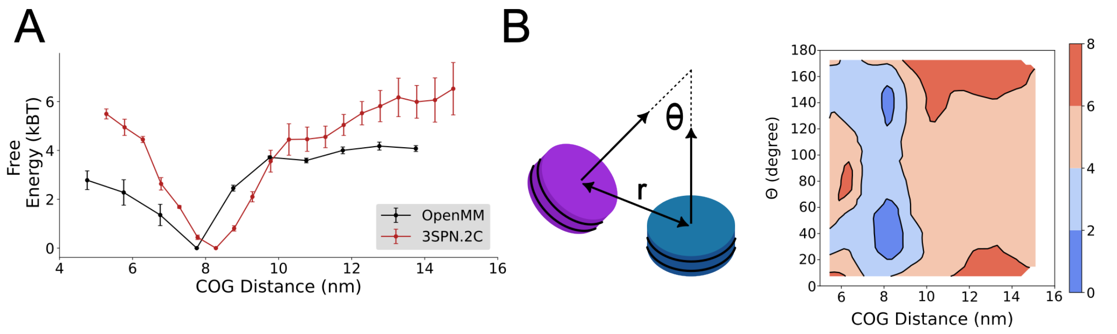


Figure S2: **Inter-nucleosome interactions for two nucleosomes.** (A) Free energy profile as a function of the geometric center distance between two nucleosomes, calculated using the explicit-ion residue-resolution model with the 3SPN.2C DNA representation (red)[S2,S5–S7] and the current model with the MRG DNA representation (black). (B) Two-dimensional free energy binding profile between two 601-sequence nucleosomes as a function of the angle $\theta$ between two nucleosome faces and the geometric center distance between the two nucleosomes.

We compared the free-energy profiles of inter-nucleosome interactions obtained from the explicit-ion version of the residue-resolution model with the 3SPN.2C DNA representation[S2,S5–S7] and from the current model with the MRG DNA representation (Figure S2). Although the two models show some differences in short-range excluded-volume repulsion, their overall free-energy profiles are semi-quantitatively similar. These differences may arise in part from the lack of DNA sequence specificity in the current MRG-based model, whereas the 3SPN.2C has been parameterized to capture sequence-dependent properties of the 601-sequence nucleosomes.[S2] Importantly, the current model captures two free-energy basins, including the most stable face-to-face configuration that stabilizes inter-nucleosome interactions, consistent with previous work using the more detailed 3SPN.2C DNA model.[S6,S8]

##### 1.1.3 12-mer Nucleosomal Array Structure

To further examine the model's accuracy in reproducing higher-order chromatin structures, we simulated equilibrated 12-mer nucleosomal arrays under three monovalent salt conditions. The simulations demonstrate that the model captures chromatin compaction across different ionic strengths and linker DNA lengths. Consistent with the DNA persistence-length calibration, a DNA bonded_scale of 0.8 quantitatively reproduces experimentally measured chromatin structure and was therefore adopted as the baseline parameter for all simulations in this manuscript.

#### 1.2 Benchmark of Model Efficiency

To evaluate the computational efficiency of the coarse-grained model, we benchmarked simulations of a 100-nucleosome system. Performance was compared across different CPU configurations and on a single NVIDIA L40S GPU to determine the optimal throughput for large-scale production runs.

Given the reduced number of degrees of freedom in the coarse-grained representation, a single 100-nucleosome simulation does not fully saturate the computational memory of a modern GPU. To maximize hardware utilization, we employed a multi-simulation strategy, running an increasing number of independent replicas concurrently on a single GPU using NVIDIA Multi-Process Service (MPS).[S10]

CPU performance was benchmarked on Intel Xeon Platinum 8462Y processors by running simulations in parallel on 8, 16, 32, and 64 CPU cores, providing a baseline for comparison with GPU performance. In the GPU benchmark, throughput increases with the number of concurrent simulations until reaching hardware saturation. At peak performance, the model achieved a maximum throughput of approximately 1,600 timesteps per second. This computational efficiency enables microsecond-scale simulations within days, allowing the extensive sampling needed to construct phase diagrams and determine the critical concentrations reported in the main text.

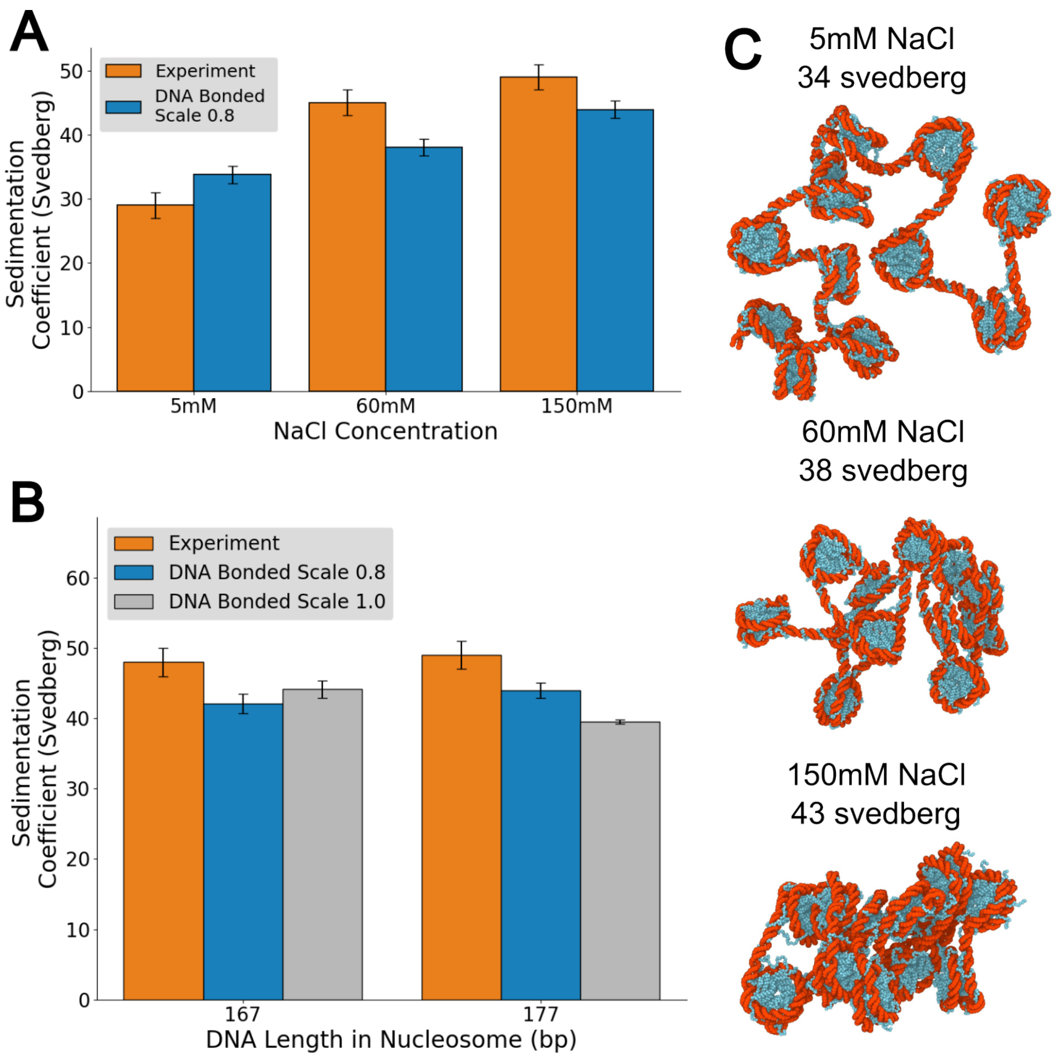


Figure S3: **Predicted sedimentation coefficients of 12-mer nucleosomal arrays under different ionic and structural conditions. (A) Salt-dependent sedimentation coefficients.** Comparison between experimental benchmarks (orange) against model-predicted sedimentation coefficients (Svedbergs, blue) for a 12-mer nucleosomal array at three monovalent salt concentrations: 5 mM, 60 mM, and 150 mM. The close quantitative agreement validates the model's ability to capture ionic-strength-dependent chromatin compaction. **(B) Influence of linker length and DNA rigidity on sedimentation.** Comparison between experimental measurements (orange) and model predictions for nucleosomal arrays with two nucleosome repeat lengths (167 bp and 177 bp). Simulations were performed using both the baseline model (bonded_scale = 0.8, blue) and a more rigid DNA model (bonded_scale = 1.0, grey), demonstrating that the model captures the sensitivity of chromatin compaction to geometric and mechanical parameters. **(C) Representative 12-mer nucleosomal array under different monovalent salt conditions.** Representative central structures were selected from the sampled ensemble at each salt concentration using the single-linkage algorithm implemented in Gromacs.[S9] Components are rendered using the main-text color scheme (DNA in red, proteins in blue).

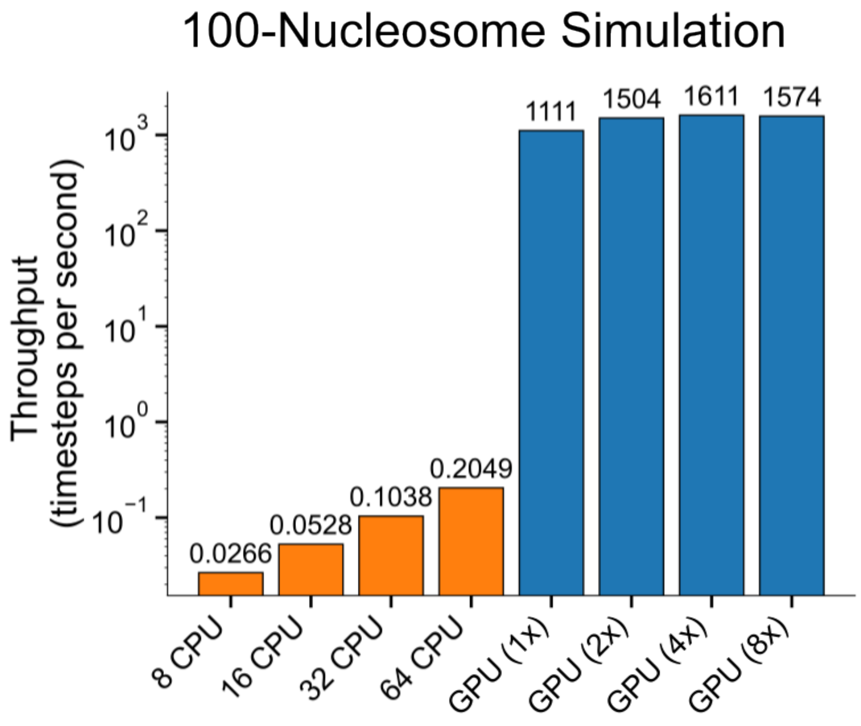


Figure S4: **Computational performance and hardware throughput benchmarking.** Average simulation throughput measured in timesteps per second (y-axis, log scale) across different hardware configurations. The first four profiles (orange) report CPU performance as a function of core allocation for a fixed simulation workload. The second set of profiles (blue) reports GPU performance for a single GPU running multiple concurrent simulation instances to determine the hardware saturation threshold. A peak multi-simulation throughput of 1,611 timesteps per second is achieved with 4 simultaneous simulations per GPU, identifying the optimal resource allocation strategy utilized for production trajectories.

#### 1.3 Determining Dense and Dilute Phases

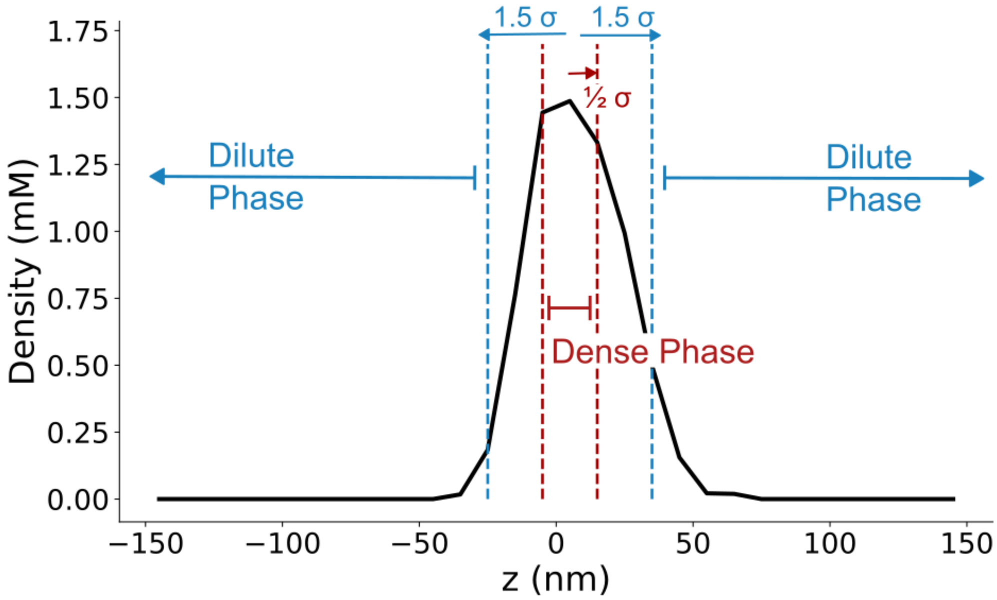


Figure S5: **Statistical definition of phase boundaries from spatial density profiles.** Representative spatial density profile along the $z$-axis of the simulation slab at 200 mM salt (black line), illustrating the procedure used to define dense and dilute phases. The system exhibits a localized condensate peak centered near the middle of the simulation box, with the density decaying to baseline values toward the box edges. Vertical dashed lines and colored arrows indicate the statistical thresholds used for phase partitioning: The dense phase (red arrows) is defined as the high-density region within $\pm 0.5\sigma$ (standard deviations) from the condensate center. The dilute phase (blue arrows) is defined as the bulk region beyond $\pm 1.5\sigma$ from the condensate center, where the density reaches a stable baseline. These spatial regions were used to extract the dense- and dilute-phase densities, $\rho_H$ and $\rho_L$, respectively, for constructing the main-text phase diagrams.

The concentrations of the dense ($\rho_{\text{high}}$) and dilute ($\rho_{\text{low}}$) phases were determined from the spatial mass-density profiles as a function of salt concentration (Figure S5). The spatial extent of the condensate was first characterized by the centroid position, $\bar{z}$, and the standard deviation, $\sigma_z$, of the density distribution:

$$\bar{z} = \frac{\sum z_i \rho_i}{\sum \rho_i}, \quad \sigma_z = \sqrt{\frac{\sum (z_i - \bar{z})^2 \rho_i}{\sum \rho_i}} \tag{S1}$$

The dense phase density was calculated by averaging the density values within the condensate core, defined as $z \in [\bar{z} - 0.5\sigma_z, \bar{z} + 0.5\sigma_z]$. Conversely, the dilute-phase density was determined by averaging the density in regions outside the condensate, defined as $z < \bar{z} - 1.5\sigma_z$ and $z > \bar{z} + 1.5\sigma_z$. Mass densities reported in $mg/mL$ were converted to molar concentrations using the nucleosome molecular weight ($M_w \approx 203.2$ kDa).

#### 1.4 Calculating Critical Concentrations

The relationship between the order parameter, $\Delta\rho = \rho_{\text{high}} - \rho_{\text{low}}$, and the salt concentration, $C$, was used to estimate the critical concentrations of the phase diagram (Figure S6). The data were fitted to an Ising-model-based expression:[S11,S12]

$$\Delta\rho = A(C_c - C)^{\beta} \tag{S2}$$

where $C_c$ is the critical salt concentration, $A$ is a fitting parameter, and $\beta = 0.325$ is the critical exponent for the three-dimensional Ising universality class. The parameters $A$ and $C_c$ were optimized using a non-linear least-squares fitting with the `scipy.optimize.curve_fit` function. The robustness of each fit was evaluated by calculating the root-mean-square error (RMSE) between the fitted and observed density differences across the corresponding range of salt concentration.

#### 1.5 Cleaning the Data for phase diagram

At certain salt concentrations, simulations did not converge to a two-phase equilibrium state and instead formed multiple discrete condensates (highlighted in red in Figures S7 and S8). These non-equilibrated states were characterized by density profiles containing multiple peaks of comparable magnitude (at 250, 260, and 275 mM) or by a large standard deviation in the dense-phase density (at 150 mM). Because these multi-droplet configurations do not adhere to a strict two-phase coexistence state, they were excluded from the binodal

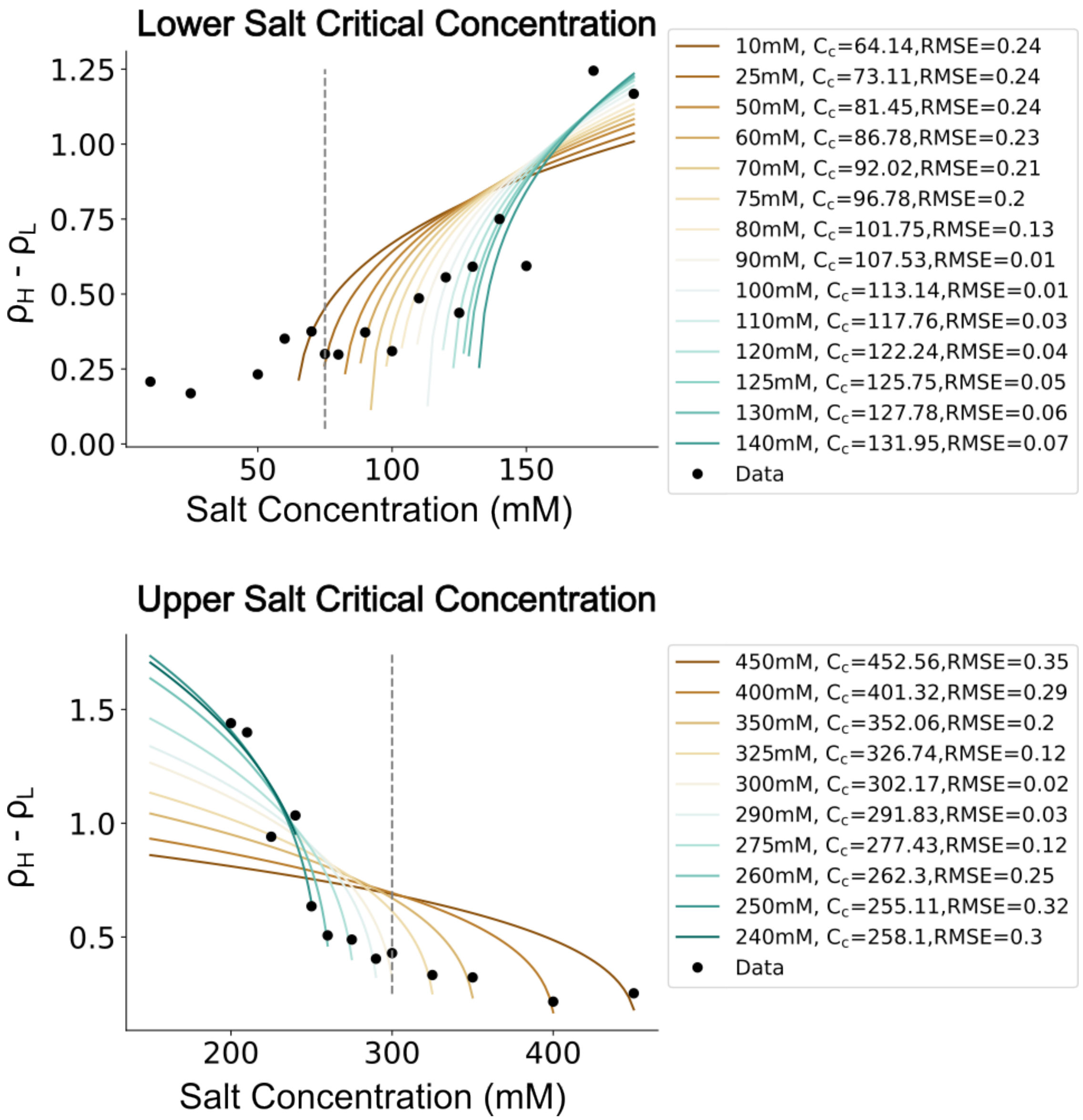


Figure S6: **Determination of lower and upper critical salt concentrations. (Top) Determination of the lower critical salt concentration ($c^*_{\text{lower}}$).** The density difference between the dense and dilute phases, $\rho_H - \rho_L$, is plotted as a function of salt concentration. Black circles represent simulation data. Colored lines (brown-to-teal gradient) represent independent fits generated by progressively truncating data points from the lower end of the range of salt concentration to isolate the lower binodal boundary. The legend lists the minimum excluded salt threshold, the resulting fitted critical concentration ($C_c$), and the corresponding root-mean-square error (RMSE). The vertical dashed grey line marks the final estimate of $c^*_{\text{lower}}$, selected based on minimization of the fitting error. **(Bottom) Determination of the upper critical salt concentration ($c^*_{\text{upper}}$).** The corresponding truncation analysis was used to identify the upper phase boundary. Colored fitting curves (brown-to-teal gradient) were generated by progressively removing high-salt data points from the fitting window. Minimization of the RMSE across these fitting windows determines the final estimate of $c^*_{\text{upper}}$ (vertical dashed grey line), above which the system transitions to a homogeneous single-phase state.

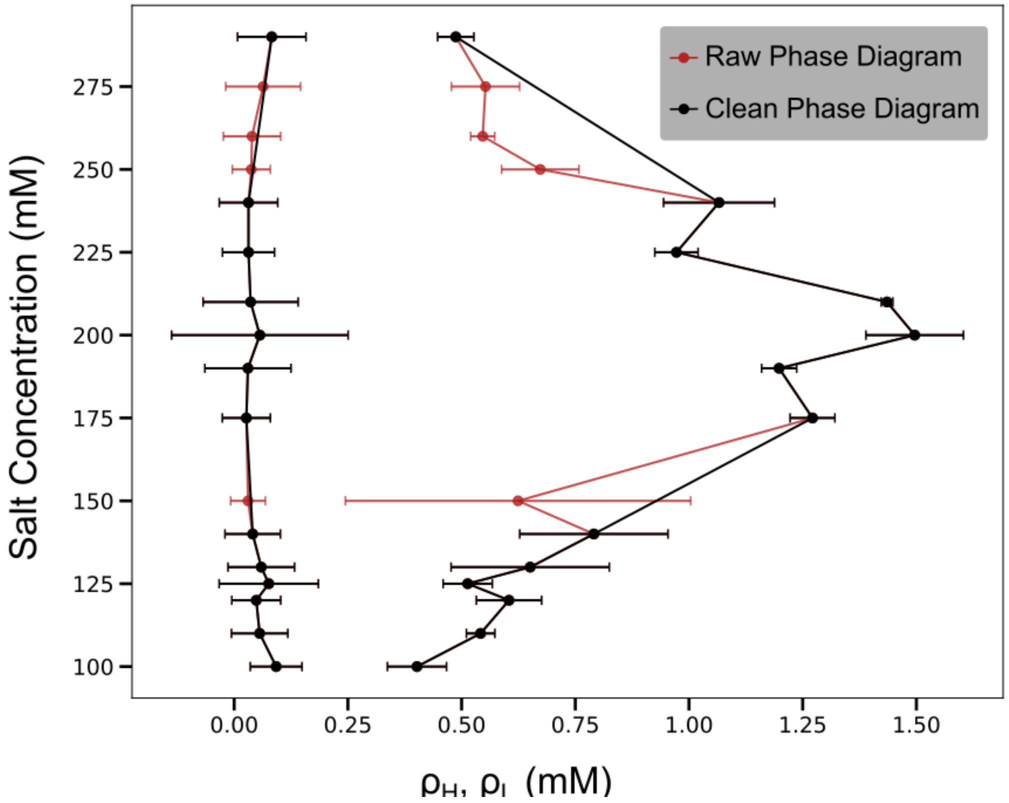


Figure S7: **Comparison of raw and cleaned phase diagrams.** Salt-dependent phase diagram showing nucleosome density (mM, x-axis) as a function of monovalent salt concentration (mM, y-axis). The red lines and data points represent the raw data extracted directly from the full set of completed trajectories. The black lines and data points represent the cleaned phase diagram presented in the main text (Figure 2). Deviations between the two curves occur at specific salt concentrations where the simulation did not reach a converged two-phase equilibrium within the simulation window. These non-equilibrium points were excluded from the main text analysis following convergence diagnostics (detailed in Fig. S8).

construction to avoid an artificial underestimation of the dense-phase density. However, these data points were retained for all other relevant analyses, including estimation of critical concentrations ($c_{\text{crit}}$).

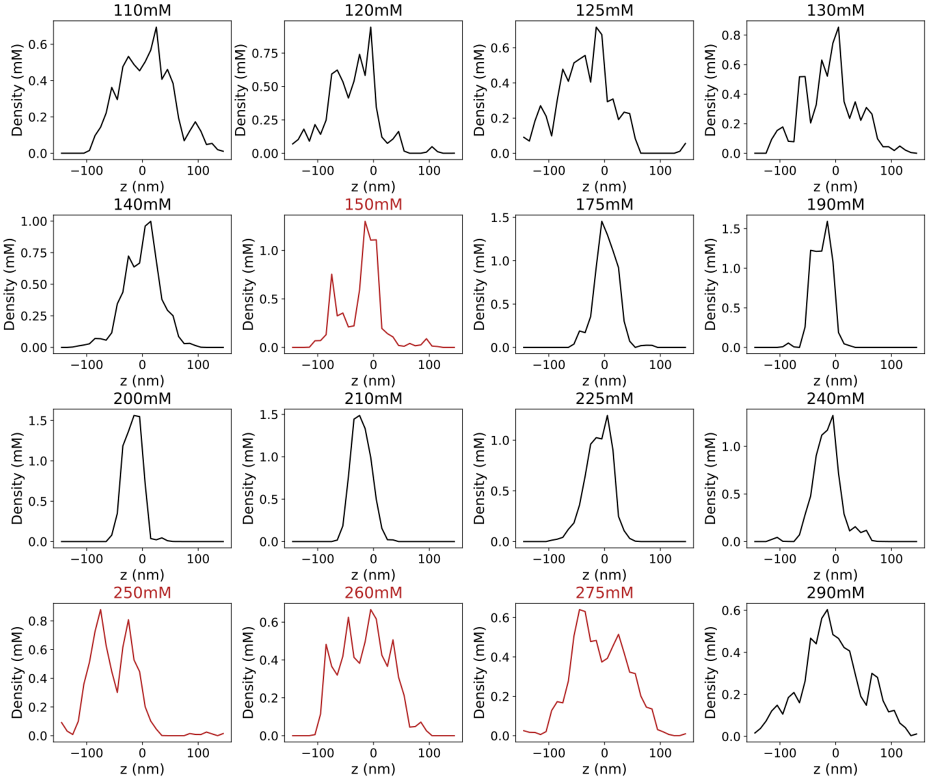


Figure S8: **Spatial density profile diagnostics for trajectory equilibration and convergence.** Spatial nucleosome density profiles calculated across individual simulation trajectories at salt concentrations within the two-phase coexistence regime. The profiles show local nucleosome density (mM) as a function of position along the simulation box. Converged trajectories (black curves) exhibit a single, well-defined central peak with low statistical variance, indicating convergence into a macroscopic phase-separated condensate. Unconverged trajectories (red curves) are characterized by large standard deviation in dense-phase density or multiple distinct peaks along the spatial axis, indicating that the system remains trapped in a multi-droplet state rather than coalescing into a single bulk condensate. These unconverged trajectories were excluded from the cleaned phase diagram (Figure S7).

#### 1.6 Additional Simulation Analyses

##### 1.6.1 Definition of Nucleosome-Nucleosome Contact Thresholds

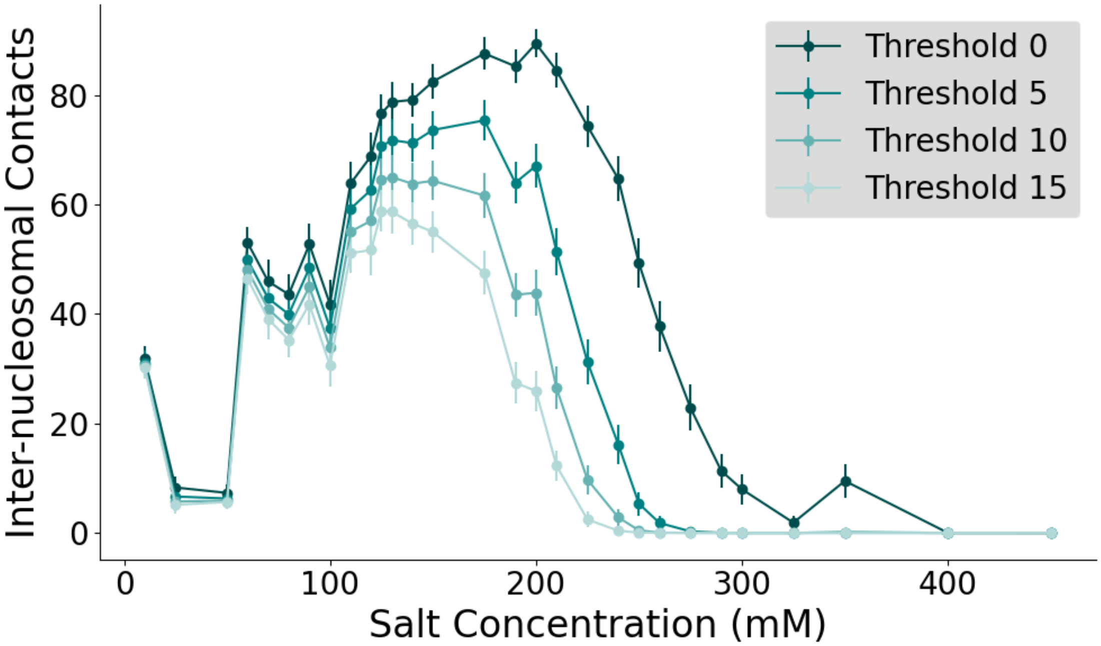


Figure S9: **Sensitivity and robustness analysis of nucleosome interaction thresholds.** Total sum of inter-nucleosomal histone-tail-DNA contacts plotted as a function of monovalent salt concentration across four distinct filtering thresholds: $\theta = 0, 5, 10,$ and 15. The interaction profiles are color-coded from dark teal ($\theta = 0$) to light teal ($\theta = 15$). While the absolute number of interaction contacts decreases as the threshold becomes more stringent, the overall salt-dependent trends and characteristic biphasic curve shapes remain consistent across all thresholds. This robustness demonstrates that the observed salt-dependent transitions are not artifacts of a specific threshold. We selected $\theta = 5$ as a balanced threshold for filtering transient contacting noise while retaining multivalent interactions in the network and contact analyses.

To quantify the degree of interactions between nucleosomes within the condensate, we analyzed the contacts between the histone tail residues of nucleosome $i$ and the DNA beads of nucleosome $j$. An inter-nucleosomal contact was recorded when the distance between a histone tail residue and a DNA nucleotide fell below a defined cutoff distance ($d_c$). Because nucleosome condensates are crowded environments, individual inter-nucleosomal contacts can represent transient thermal fluctuations rather than stable structural interactions. To distinguish stable inter-nucleosomal contacts from this transient noise, we introduced a contact threshold, $\theta$, which defines the minimal number of amino-acid-nucleotide contacts required to classify two nucleosomes as being in contact. We performed a sensitivity analysis by calculating interaction profiles across a range of thresholds ($\theta = 0, 5, 10,$ and 15) (Figure S9). While the absolute number of inter-nucleosomal contacts decreases with increasing threshold, the overall shape and relative trends across the salt concentrations remain largely invariant. The preservation of the salt-dependent profile across different thresholds indicates that our findings are robust and not the artifact of a specific cutoff choice. We therefore selected a non-zero threshold ($\theta = 5$) for all the analyses presented in the main text. This threshold filters out stochastic, short-lived interactions and focuses the analysis on the stable, multivalent "sticking" events that stabilize condensates.

##### 1.6.2 Spatiotemporal Analysis of Nucleosome Interaction Networks

To elucidate the microscopic connectivity driving the system's macroscopic phase separation, we analyzed the nucleosome interaction networks at three representative salt concentrations: 10 mM (low salt), 200 mM (intermediate/physiological salt), and 400 mM (high salt) (Figure S10).

**Time-averaged contact probability:** The top panels of Figure S10 characterize contact occupancy, defined as the fraction of simulation time during which two nucleosomes remain in contact ($0 \leq P \leq 1$). At 10 mM, the contact map shows a sparse distribution of high-intensity spots, indicating a regime dominated by a small number of stable, long-lived contacts. These contacts likely represent small clusters of strongly interacting nucleosomes. At 200 mM, the contact map shows a broad distribution of weaker contacts, representing a highly dynamic network where nucleosomes form many transient and constantly exchanging interactions. This is characteristic of a liquid-like condensate where molecular "stickers" frequently exchange partners.[S13] At 400 mM, the absence of significant contact occupancy indicates a fully dissociated state where electrostatic screening and thermal fluctuations prevent stable inter-nucleosome association.

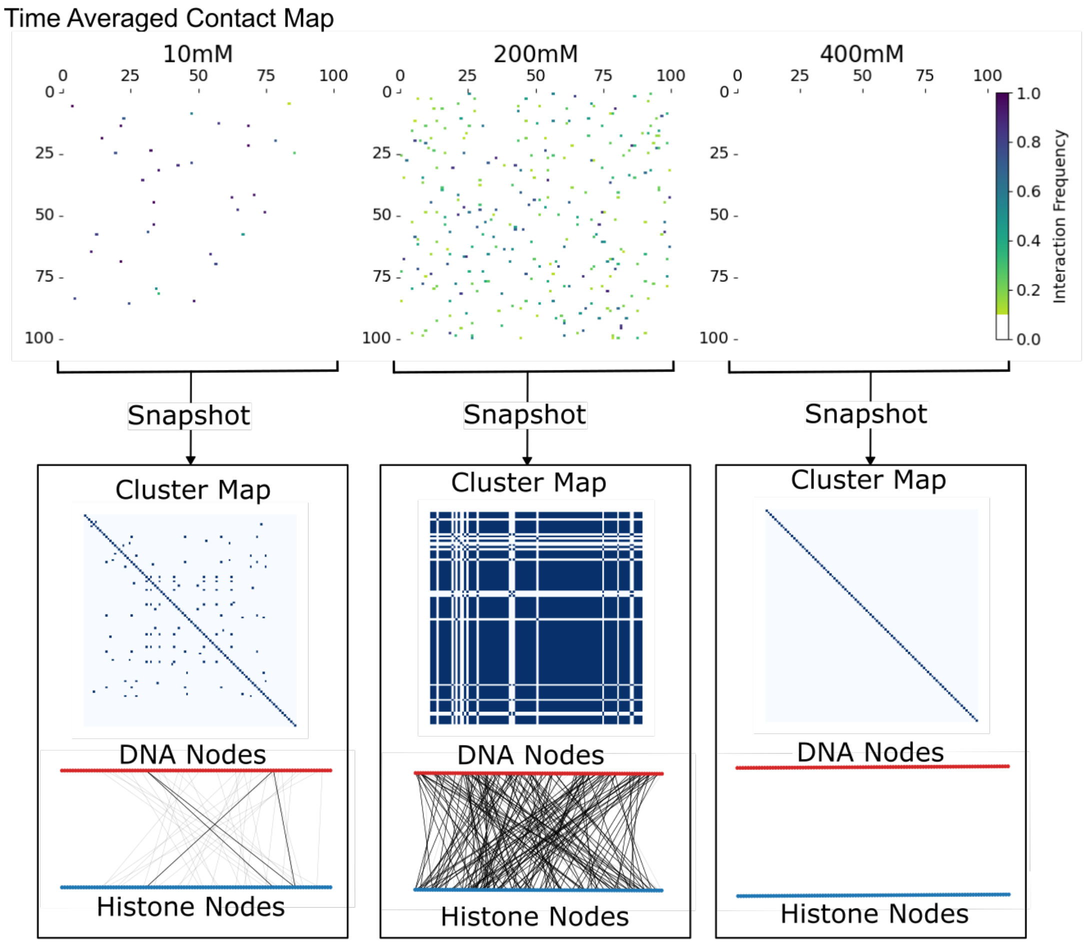


Figure S10: **Salt-dependent evolution of chromatin networks.** (Top) Time-averaged contact maps showing the transition from localized, stable nucleosome interactions at 10 mM salt to broad, dynamic network formation at 200 mM salt, followed by dissociation into isolated nucleosomes at 400 mM salt. (Middle) Instantaneous binary cluster maps at different salt concentrations. (Bottom) Node-link diagrams of nucleosome interaction networks, where bold edges indicate the largest connected component (LCC). The larger LCC at 200 mM ($N > 80$) relative to that at 10 mM ($N \approx 5$) highlights the salt-induced transition from isolated microphase clusters to a globally connected condensate network.

**Snapshot network topologies (Cluster and Node Maps):** To capture the instantaneous connectivity of the system, we analyzed individual simulation frames using binary cluster maps and node-link diagrams. The binary cluster maps (Figure S10 middle panels) represent the system as discrete groups of connected nucleosomes. At 10 mM, nucleosomes form small, isolated clusters but do not assemble into a system-wide network. At 200 mM, the system is dominated by large, system-spanning clusters, indicating the formation of a percolated network characteristic of a macroscopic condensate. At 400 mM, no clusters are detected, consistent with a dilute, dissociated phase.

**Node-Link connectivity and largest cluster analysis:** In the node-link diagrams (Figure S10 bottom panels), nodes represent individual nucleosomes (Red: DNA, Blue: Protein), while edges (grey lines) represent all instantaneous contacts. The bold overlay identifies the largest connected component (LCC). At 10 mM, the LCC is small (containing only $\sim 5$ connections), indicating limited network connectivity. At 200 mM, the LCC is significantly larger (containing $> 80$ connections), demonstrating that nearly the entire system is integrated into a single, cohesive network. At 400 mM, the absence of edges confirms the loss of stable inter-nucleosome contacts under high-salt conditions.

##### 1.6.3 Mechanical and Chemical Modulation of Condensate Architecture

**DNA unwrapping propensity as a function of bending rigidity:** To investigate the mechanical origin of the phase transitions, we quantified the degree of DNA unwrapping from the histone core (Figure S11). We observe a clear correlation between DNA bending rigidity, controlled by the bonded scale parameter, and the onset of DNA unwrapping. Arrays with higher rigidity exhibit significantly greater DNA unwrapping across ionic concentrations than more flexible arrays. Increased rigidity raises the energetic cost of maintaining DNA in its wrapped conformation around the histone core. This mechanical tension facilitates the release of DNA segments, thereby increasing the availability of both DNA and histone surfaces for inter-nucleosomal interactions. This result indicates that DNA rigidity lowers

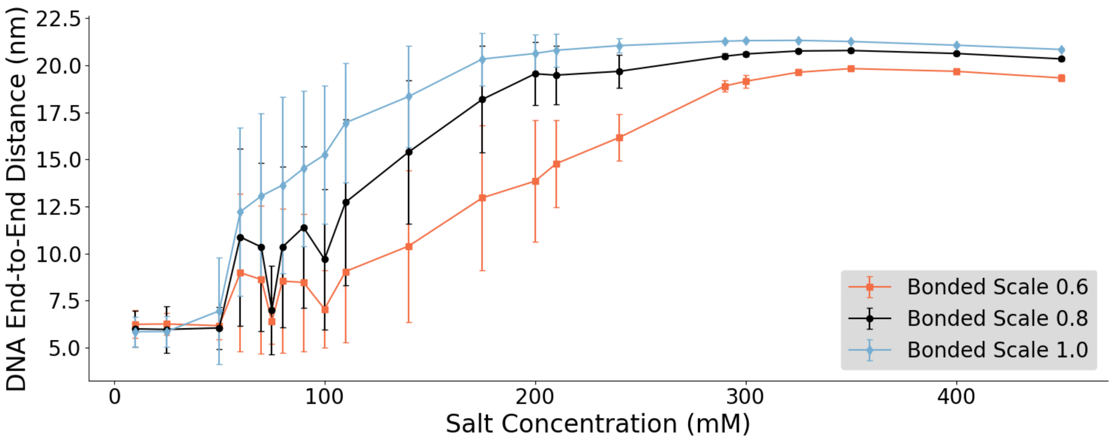


Figure S11: **DNA unwrapping propensity as a function of polymer bending rigidity.** Average end-to-end distances of nucleosomal DNA plotted as a function of monovalent salt concentration across different DNA bending rigidities. The profiles follow the main text color scheme: bonded scale = 0.6 (orange), bonded scale = 0.8 (unmodified baseline, black), and bonded scale = 1.0 (light blue). Stiffer DNA arrays exhibit a greater propensity for spontaneous unwrapping across ionic concentrations than more flexible arrays. This increased mechanical tension lowers the energetic barrier for the release of wrapped DNA segments, expands the available surface area for inter-nucleosomal interactions, and provides a structural basis for the denser condensate.

the salt-dependent barrier for the conformational changes necessary to drive condensation.

**Comparative density profiles: unmodified versus acetylated arrays** To connect our physical model to epigenetic regulation, we compared the local density distributions of unmodified nucleosome condensates with those containing histone tail lysine acetylations (Figure S12). Acetylated lysines were modeled by neutralizing the corresponding positive charges. Unmodified nucleosomes exhibit a sharply peaked density distribution within the condensate core, indicating compact organization stabilized by strong electrostatic attractions between histone tails and DNA. In contrast, acetylated nucleosome condensates display broader density profiles with low peak density, consistent with a less compact organization. These results are consistent with the findings of Bascom and Schlick (2018),[S14] which show that acetylation-induced reduction in tail-DNA affinity leads to a more open chromatin state. Interestingly, this chemical loosening resembles certain aspects of the mechanical loosening observed with high-rigidity DNA, suggesting that both chemical modifications and mechan-

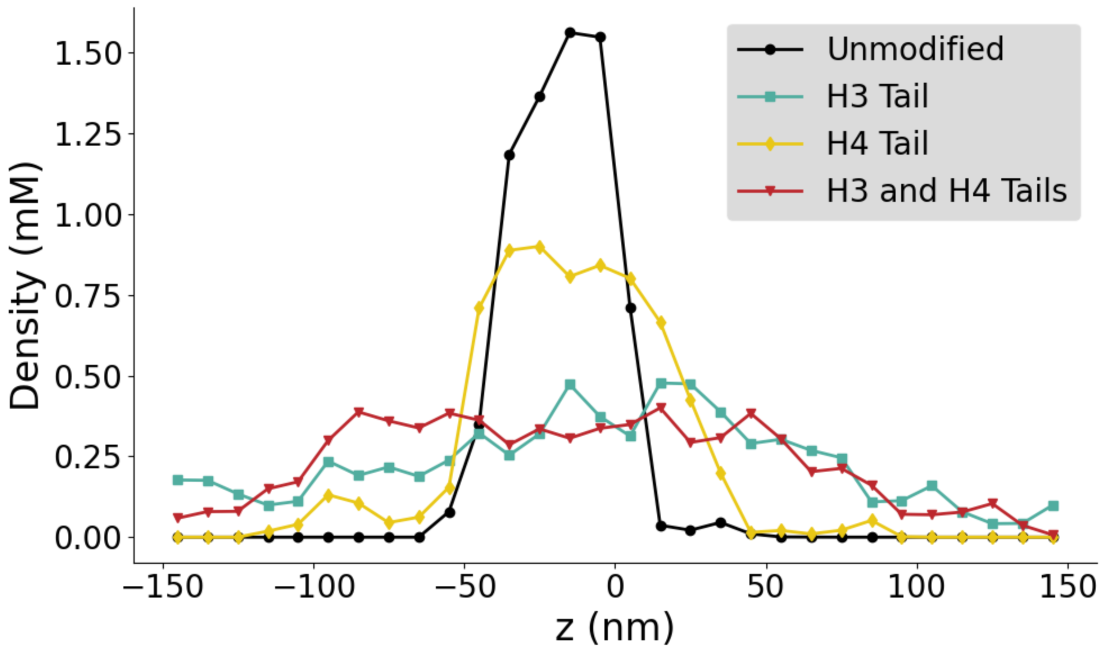


Figure S12: **Spatial density profiles of chromatin condensates across tail acetylation states.** Spatial nucleosome density profiles along the $z$-axis of the simulation slab at 200 mM salt for different histone tail acetylation states. The profiles follow the main text color and labeling scheme: unmodified nucleosomes (black), H3-tail-acetylated nucleosomes (teal), H4-tail-acetylated nucleosomes (yellow), and dual H3/H4-tail-acetylated nucleosomes (red). The unmodified system exhibits a sharp and compact central density peak. Histone-tail acetylation progressively broadens the spatial density profile and reduces the peak density, with the combined H3/H4 acetylation producing the most expanded and diffuse internal condensate architecture.

ical properties can be used to tune the density and accessibility of chromatin condensates in cells.

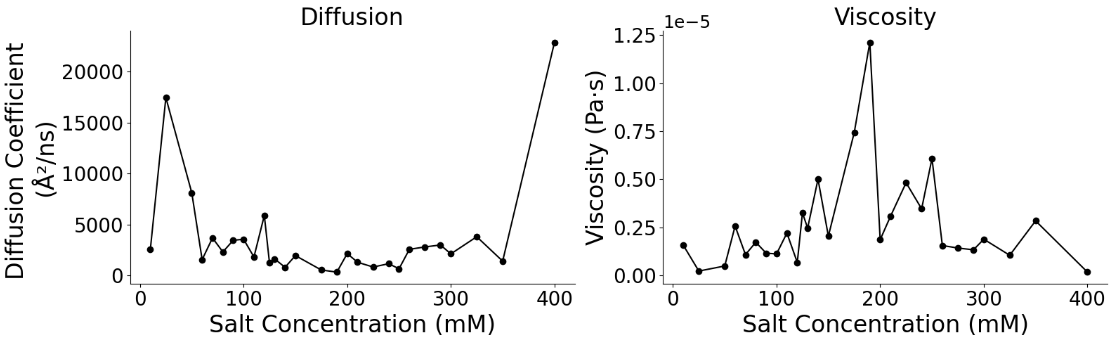


Figure S13: **Dynamic and rheological profiles of the chromatin condensate across the two-phase coexistence regime. (Left) Salt-dependent diffusion coefficients ($D$).** Calculated nucleosome diffusion coefficients, reported in the unit of $\text{Å}^2$/ns, for the wild-type nucleosome system (bonded scale = 0.8) across the evaluated range of monovalent salt concentrations (10 mM to 400 mM). The sharp increase in diffusion near the low- and high-salt boundaries (25 mM and 400 mM) reflect accelerated nucleosome dynamics as the system approaches its lower and upper critical concentrations ($c^*$), where the condensate dissolves into homogeneous single-phase state. **(Right) Salt-dependent condensate viscosity ($\eta$).** Effective viscosity (Pa·s) across the same salt-concentration range. The viscosity shows an approximately inverse relationship with diffusion, featuring a peak in the intermediate physiological salt regime (150–200 mM). This peak coincides with the maximum dense-phase density and strongest inter-nucleosomal contact connectivity of the wild-type nucleosomal network.

(S1) Hagerman, P. J. Flexibility of DNA. *Annu. Rev. Biophys. Biophys. Chem.* **1988**, *17*, 265–286.

(S2) Freeman, G. S.; Hinckley, D. M.; Lequieu, J. P.; Whitmer, J. K.; De Pablo, J. J. Coarse-grained modeling of DNA curvature. *The Journal of Chemical Physics* **2014**, *141*, 165103.

(S3) Geggier, S.; Vologodskii, A. Sequence dependence of DNA bending rigidity. *Proc Natl Acad Sci U S A* **2010**, *107*, 15421–15426.

(S4) Eastman, P.; Galvelis, R.; Peláez, R. P.; Abreu, C. R. A.; Farr, S. E.; Gallicchio, E.; Gorenko, A.; Henry, M. M.; Hu, F.; Huang, J.; Krämer, A.; Michel, J.; Mitchell, J. A.; Pande, V. S.; Rodrigues, J. P.; Rodriguez-Guerra, J.; Simmonett, A. C.; Singh, S.; Swails, J.; Turner, P.; Wang, Y.; Zhang, I.; Chodera, J. D.; De Fabritiis, G.; Markland, T. E. OpenMM 8: Molecular Dynamics Simulation with Machine Learning Potentials. *J. Phys. Chem. B* **2024**, *128*, 109–116.

(S5) Freeman, G. S.; Hinckley, D. M.; De Pablo, J. J. A coarse-grain three-site-per-nucleotide model for DNA with explicit ions. *The Journal of Chemical Physics* **2011**, *135*, 165104.

(S6) Lin, X.; Zhang, B. Explicit ion modeling predicts physicochemical interactions for chromatin organization. *eLife* **2024**, *12*, RP90073.

(S7) Li, R.; Lin, X. Connected Chromatin Amplifies Acetylation-Modulated Nucleosome Interactions. *Biochemistry* **2025**, *64*, 1222–1232.

(S8) Moller, J.; Lequieu, J.; De Pablo, J. J. The Free Energy Landscape of Internucleosome Interactions and Its Relation to Chromatin Fiber Structure. *ACS Cent. Sci.* **2019**, *5*, 341–348.

(S9) Abraham, M. J.; Murtola, T.; Schulz, R.; Páll, S.; Smith, J. C.; Hess, B.; Lindahl, E. GROMACS: High performance molecular simulations through multi-level parallelism from laptops to supercomputers. *SoftwareX* **2015**, *1-2*, 19–25.

(S10) Hsu, D.; Clark, D.; Paulsen, J. Maximizing OpenMM Molecular Dynamics Throughput with NVIDIA Multi-Process Service. NVIDIA Technical Blog, 2025; https://developer.nvidia.com/blog/maximizing-openmm-molecular-dynamics-throughput-with-nvidia-multi-process-service/ Accessed: 2026-07-03.

(S11) Dignon, G. L.; Zheng, W.; Kim, Y. C.; Best, R. B.; Mittal, J. Sequence determinants of protein phase behavior from a coarse-grained model. *PLoS Comput Biol* **2018**, *14*, e1005941.

(S12) Farr, S. E.; Woods, E. J.; Joseph, J. A.; Garaizar, A.; Collepardo-Guevara, R. Nucleosome plasticity is a critical element of chromatin liquid–liquid phase separation and multivalent nucleosome interactions. *Nat Commun* **2021**, *12*, 2883.

(S13) Wang, J.; Choi, J.-M.; Holehouse, A. S.; Lee, H. O.; Zhang, X.; Jahnel, M.; Maharana, S.; Lemaitre, R.; Pozniakovsky, A.; Drechsel, D.; Poser, I.; Pappu, R. V.; Alberti, S.; Hyman, A. A. A Molecular Grammar Governing the Driving Forces for Phase Separation of Prion-like RNA Binding Proteins. *Cell* **2018**, *174*, 688–699.e16.

(S14) Bascom, G. D.; Schlick, T. Chromatin Fiber Folding Directed by Cooperative Histone Tail Acetylation and Linker Histone Binding. *Biophys J* **2018**, *114*, 2376–2385.
